# Circuit regulation along an activity continuum shapes flexible and stable representations during learning

**DOI:** 10.64898/2026.10.28.762663

**Authors:** Daniel Yusufi, Yan Ma, Lujia Chen, Garret Wang, Taylor Malone, Kai Zhang, June Hoan Kim, Farid Shahid, Kyle Cekada, Ellane Barcelone, Yi Gu

**Author notes:** Corresponding author: Yi Gu.

## Abstract

Learning requires neural representations to flexibly incorporate new information while progressively stabilizing to support enduring memories. How neural circuits balance these demands remains unclear. Here, we identify complementary mechanisms supporting representational flexibility and stability in the medial entorhinal cortex (MEC), a key component of the cognitive map for memory-guided navigation. Longitudinal calcium imaging in mice learning novel virtual environments revealed a persistent axis of excitatory neuronal activity associated with distinct representational dynamics. Low-activity neurons exhibited intermittent activation that varied with learning stages and greater learning-dependent representational refinement, flexibly shifting their encoding from landmarks toward reward locations. By contrast, high-activity neurons exhibited more persistent activation and maintained more stable spatial representations. Dopaminergic pre-reward ramping was associated with the emergence of reward encoding in low-activity neurons, whereas interactions with inhibitory interneurons contributed to the stability of high-activity neurons. Notably, interneurons exhibited activity changes opposing those of excitatory neurons during learning, and their transient activation upon novelty exposure delayed excitatory representational remodeling, stabilization, and behavioral learning, suggesting that inhibitory circuits gate learning-dependent representational dynamics. Together, these findings reveal how activity-dependent excitatory neuronal organization and circuit-level regulation enable complementary forms of representational flexibility and stability to coexist, facilitating the incorporation of novel experiences into stable, context-specific cognitive maps that support learning.

## Introduction

Learning requires neural representations to balance two competing demands: stability across experience and flexibility in response to newly acquired information. This stability-flexibility dilemma poses a fundamental challenge: excessive stability can impede new learning, whereas excessive flexibility can disrupt established representations^1–3^. How neural circuits balance these demands during memory formation remains a central question in learning and memory.

This challenge is particularly evident during spatial learning in a stable environment (Fig. 1a). Although the physical features of an environment may remain unchanged, their behavioral significance can evolve with experience. Salient landmarks or boundaries may initially dominate neural representations, but as animals learn task structure, behaviorally relevant features such as reward locations may become increasingly important. Thus, spatial learning requires the brain to preserve a stable representation of the environment while flexibly incorporating newly acquired knowledge, making spatial learning a valuable framework for investigating how neural circuits balance representational stability with experience-dependent flexibility.

**Figure 1.**
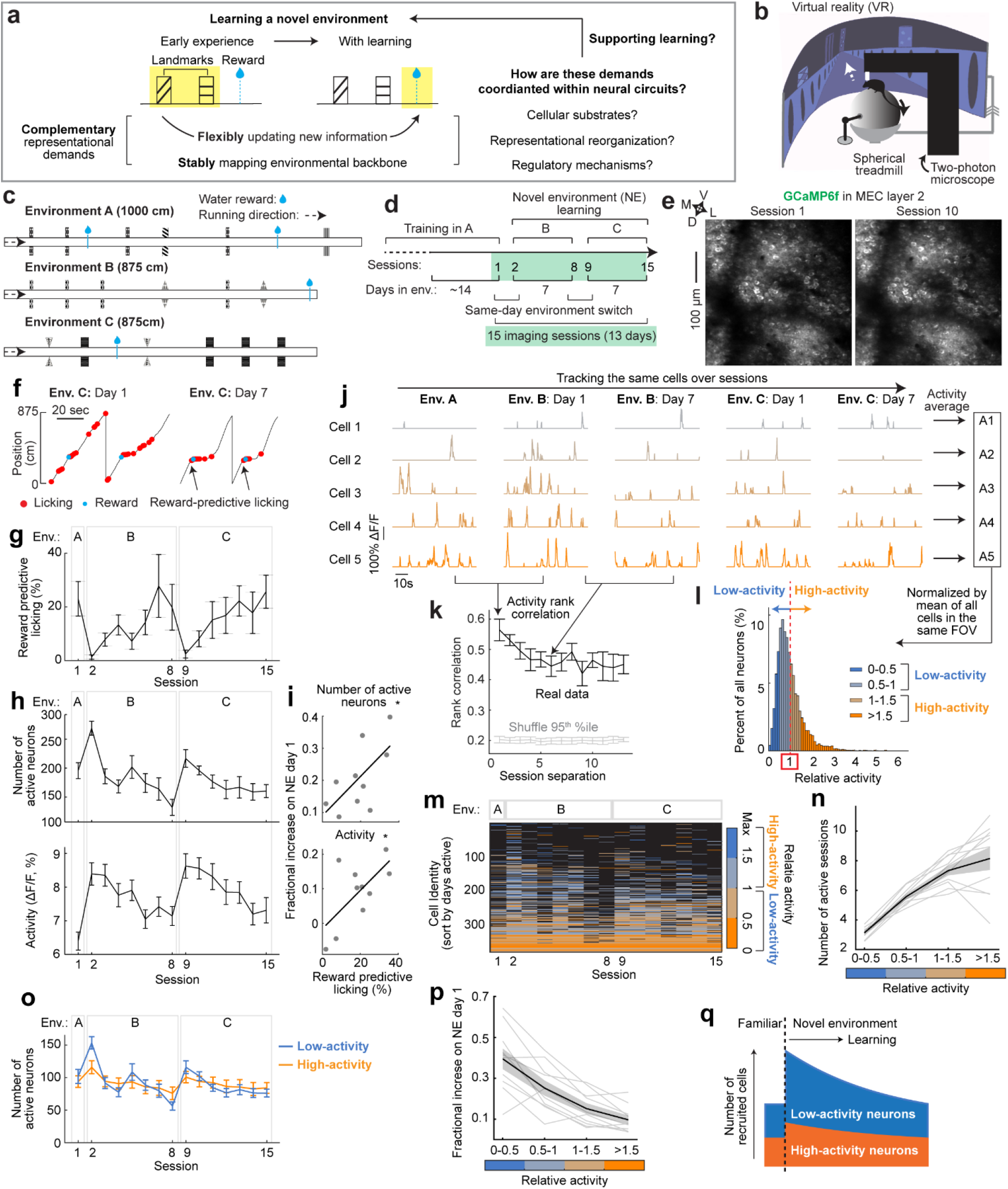
Neuronal activity level organizes recruitment dynamics during novel environment learning. **a.** Schematic illustrating the complementary representational demands that the current study aims to address. **b.** Schematic of the head-fixed virtual reality (VR) and two-photon setup. **c.** Familiar environment A (1000 cm) and novel environments B and C (875 cm each). Icons indicate visual landmarks and the water symbol marks the reward locations. **d.** Experimental timeline: Mice were imaged in familiar environment A and subsequently for 7 days each in novel environments B and C. Sessions 1 and 2 occurred on the same day, as did sessions 8 and 9, resulting in 15 imaging sessions across 13 days. **e.** Example two-photon images of GCaMP6f-expressing MEC layer 2 neurons tracked through two imaging days. **f.** Example behavior on days 1 and 7 in environment C (Env. C). Red and blue dots indicate licks and reward delivery, respectively. **g.** Reward-predictive licking across sessions. **h.** Number of active neurons (top) and mean activity (bottom) of neurons detected across all sessions. **i.** Pearson correlation between novelty-associated increases in the number of active neurons (top) or neural activity (bottom), as defined in **h**, and reward-predictive licking during late-learning (days 4-7) in the subsequent environment. **j.** Example significant calcium transients from five neurons tracked across environments and ordered by their mean activity across sessions. **k.** Spearman correlation of neural activity rank as a function of session difference. Black, real data; gray, 95^th^ percentile of shuffled correlations obtained by permuting neural activity ranks. **l.** Distribution of relative activity, defined as each neuron’s mean activity across all sessions in which it was detected normalized by the mean of these neuron-wise averages within the same FOV. Colors indicate four relative activity groups. **m.** Visualization of detection of longitudinally tracked neurons across sessions. Neurons are colored by relative activity and sorted by the number of sessions in which they were active. Matrix is from one representative FOV. **n.** Number of sessions in which neurons were active as a function of relative activity. Thin lines indicate individual FOVs and black indicates the mean. **o.** Number of active low-activity and high-activity neurons across all sessions. **p.** Fractional increase in active neurons on the first day of novel environment exposure as a function of relative activity. **q.** Summary schematic demonstrating preferential recruitment of low-activity neurons during novelty and their further refinement with learning, whereas high-activity neurons show more persistent activation. Data are mean ± s.e.m. unless otherwise indicated. *, *P* ≤ 0.05, **, P ≤ 0.01, ***, *P* ≤ 0.001; n.s., *P* > 0.05. The dataset comprised 10 FOVs from 5 mice (2 FOVs per mouse). i, *n* = 10 (5 mouse × 2 learning-environment, data were averaged across FOV from the same mice).

The hippocampal-entorhinal system is central to spatial learning and is well positioned to support this balance. It contains neurons that are engaged during spatial navigation, including place cells in the hippocampus^4^, grid cells in the medial entorhinal cortex (MEC)^5^, and neurons encoding head direction^6^, boundaries^7^, landmarks^8–11^, and speed^12,13^. Hippocampal representations tend to be flexible, as place-cell recruitment and spatial firing patterns can change substantially across environments^14–16^ and even within the same environment^13,17^. In contrast, MEC representations often exhibit greater stability, with similar neural ensembles recruited within and across environments^18^ and stable activity patterns developing with experience^18,19^. Coupling between grid cells also remains coherent despite changes to the environments and experience^18,20–22^. Although these past findings suggest largely complementary roles for the hippocampus and MEC in flexibility and stability, this division is unlikely to be absolute. Notably, the MEC can support spatial learning even after hippocampal lesions^23^, and its activity is strongly influenced by environmental features^24–28^, landmarks^29–31^, and reward^18,32,33^. Thus, flexible and stable spatial signals may coexist within the same MEC network, allowing newly acquired information to be locally integrated within a stable spatial framework. How these signals are distributed across MEC neurons, how they are shaped with learning and regulated mechanistically, and how they support the formation of robust memories remain unclear (Fig. 1a).

To address these questions, we performed longitudinal two-photon calcium imaging of MEC excitatory neurons as mice learned to navigate two novel virtual linear environments. We identified neuronal populations spanning a range of activity levels that differentially supported representational flexibility and stability under dopaminergic and inhibitory regulation, respectively. Interneurons also reduced their activity upon novelty exposure, potentially gating learning-dependent dynamics. Consistent with this, preventing this reduction delayed the emergence of flexible and stable representational dynamics and slowed behavioral learning. Collectively, these findings reveal how activity-level organization and circuit regulation together enable flexible and stable representations to coexist within the MEC to support spatial learning.

## Results

### Neuronal activity level organizes recruitment dynamics during novel environment learning

To investigate how the MEC coordinates flexible and stable representations during learning, we longitudinally imaged the calcium activity of MEC layer 2 excitatory neurons as mice learned to navigate unidirectionally along linear virtual reality (VR) environments for water rewards (Fig. 1b). Mice were first familiarized with environment A (familiar environment, FE), and then further exposed consecutively to two novel environments (NE), B and C, with 7 daily learning sessions in each environment. The environments differed in track length, wall patterns, landmark identities and arrangement, and reward locations (Figs. 1c, d). Using GP5.3 mice stably expressing the genetically encoded calcium indicator GCaMP6f in MEC layer 2 excitatory neurons^34–36^, we tracked the same neuronal populations across days and environments at cellular resolution (Fig. 1e). In both environments, mice showed progressively increased reward-predictive licking, indicating learning of the reward locations (Fig. 1f, g).

To characterize neural dynamics during learning, we defined active neurons as those with detectable calcium activity in a given session and quantified each neuron’s activity as its mean calcium response over time. This measure was mostly positively correlated with the frequency, amplitude, and duration of significant calcium transients (Fig. S1a), supporting its use to reflect neuronal response.

Learning was accompanied by pronounced changes in MEC population activity. Novelty rapidly increased MEC population activity, reflected by increases in both the number of active neurons and the activity of neurons active across all sessions (Fig. 1h). Both measures gradually declined with experience. These transient activity increases were not attributable to faster running (Fig. S1b). Notably, the novelty-associated increases in active neuron number and activity on NE day 1, relative to the preceding FE, were correlated with reward-predictive licking during late learning in the same NE (Fig. 1i), linking stronger novelty responses to better subsequent learning.

Despite these population-level changes, neurons maintained their relative activity levels across different learning stages and environments (Fig. 1j). Indeed, activity rank correlations of simultaneously imaged neurons across session pairs exceeded the 95^th^ percentile of shuffled controls in which neuronal identities were permuted (Fig. 1k). Based on this persistent activity organization, we defined each neuron’s “relative activity” as its averaged activity across sessions in which it was active, normalized by the mean of the simultaneously imaged population within each field of view (FOV). Relative activities formed a skewed, continuous distribution, resembling the distributions reported from the hippocampal-entorhinal circuit^37^ (Fig. 1l). For subsequent analysis, neurons with relative activities below or above the population mean were referred to as low- and high-activity, respectively. Cross-validation using activity from other sessions, environments, or learning days correctly predicted these classifications with approximately 85% accuracy (Fig. S1d), supporting the robustness of this categorization and the persistence of relative activity across sessions (Fig. 1k). Where applicable, we also further subdivided neurons into four relative-activity categories (0-0.5, 0.5-1, 1-1.5, and >1.5) to assess continuous relationships between activity and neuronal properties (Fig. 1l).

We next asked whether relative activity was associated with population recruitment across learning and environments. When sorting neurons by the number of sessions they were active, neurons with higher relative activity tended to be more persistently active, whereas those with lower relative activity were recruited more intermittently (Fig. 1m). Consistent with this pattern, the average number of sessions in which neurons were active increased progressively with relative activity (Fig. 1n). Notably, the novelty-associated increase in active neurons was more pronounced among the lower-activity populations (Figs. 1o, p). Further analyses indicated that this recruitment pattern was more closely associated with learning stage than with environmental identity, particularly among low-activity neurons (Figs. S2a-e). These results point toward an activity-stratified organization of neuronal recruitment across learning stages, with high-activity neurons exhibiting more persistent participation and low-activity neurons showing greater intermittent recruitment, particularly upon exposure to a novel environment (Fig. 1q).

Collectively, our findings reveal a persistent continuum of MEC neuronal activity despite dramatic learning-associated changes in population activity across environments. The novelty-associated increase in population activity was associated with subsequent learning and preferentially recruited low-activity neurons, which showed more intermittent, learning-stage-dependent recruitment than high-activity neurons. These findings implicate low-activity neurons in successful spatial learning and suggest differential contributions of low- and high-activity neurons to learning. Additionally, low- and high- activity neurons were anatomically segregated (Figs. S2f-h) and differed in their composition of major putative molecular cell types, including pyramidal and stellate cells (Figs. S2h and j), further supporting activity-dependent functional heterogeneity in the MEC circuit.

Such heterogeneity has been reported in the hippocampus, where low-activity neurons exhibited greater learning-dependent plasticity^38^ and greater representational drift in familiar environments^13^. However, whether the relationship between neural activity and representational flexibility extends to other brain regions, such as the MEC, remains unclear. It is also unknown whether these differences reflect variation in nonspecific representational stability or correspond to structured spatial coding regimes during learning. We therefore next examined whether activity is related to the spatial representations MEC neurons encode during novel environment learning.

### Neural activity organizes structured spatial representations with varying flexibility and stability during novel environment learning

To examine how relative activity relates to spatial encoding during learning, we compared the spatial activity of low- and high-activity neurons. Within sessions, low- activity neurons showed sparser spatial activity than high-activity neurons (Fig. 2a), with lower spatial field coverage, defined as the proportion of spatial bins belonging to fields comprising consecutive bins with significant calcium responses (Fig. 2b). Across learning, low-activity neurons exhibited reduced cross-session stability (Fig. 2c), reflected by weaker activity correlations between session pairs (Fig. 2d). The proportions of putative grid cells^5,39^ cue cells^8^, and speed cells^12^ also differed between activity groups, with more unclassified cells among low-activity neurons (Fig. S3d). These differences largely persisted after matching calcium events of high- and low-activity neurons in frequency, duration, and amplitude (“activity-matched”; Figs. S3a-d). Together, these observations revealed relatively rich and stable spatial representations in high-activity neurons, while raising the question of whether low-activity neurons are simply weakly tuned cells with unstable activity and limited spatial information.

**Figure 2.**
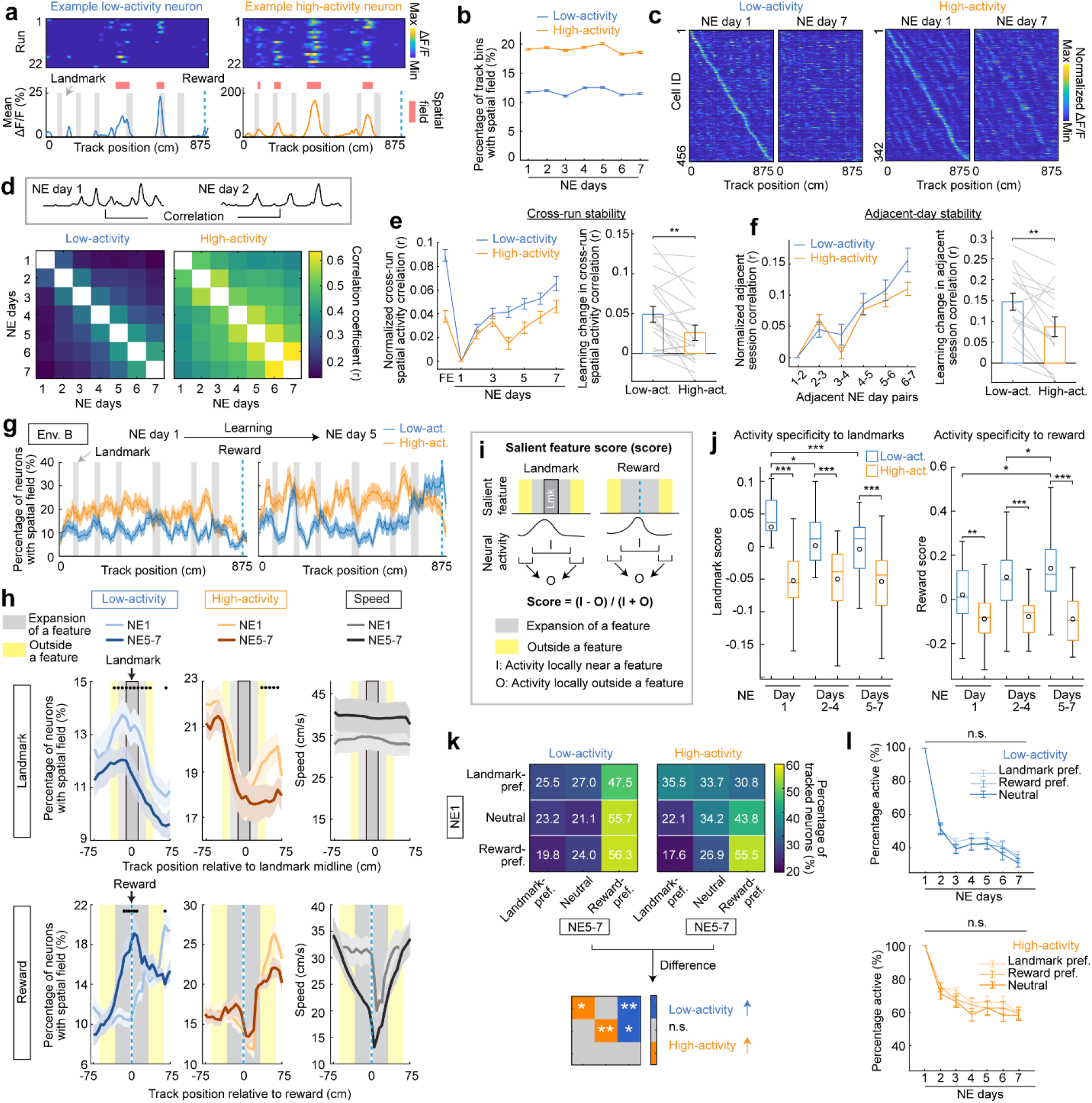
Activity organizes flexible and stable spatial representations during novel- environment learning. **a.** Example spatial activity from a low-activity and high-activity neuron. Top, run-by-run spatial activity; bottom, mean spatial activity. Red shading indicates spatial fields. **b.** Percentage of track spanned by spatial fields for low-activity and high-activity neurons across novel-environment learning. Neurons were pooled across FOVs and learning environments. **c.** Run-averaged spatial activity of low-activity and high-activity neurons tracked on NE days 1 and 7 in environment B, with neurons ordered by their NE day 1 activity peak location. Neurons were pooled across FOVs. **d.** Top, schematic of the spatial activity correlation analysis. Bottom, mean pairwise spatial correlations across the 7 learning days for low-activity and high-activity neurons. Correlations were pooled across neurons and environments. **e.** Cross-run stability across learning, with NE day 1 correlations subtracted (left), learning associated change from NE day 1 to late learning (right). Gray lines (right) indicate individual FOV × learning-environment observations (n = 20 from 10 FOVs × 2 environments; paired two-sided t-tests). **f.** Spatial activity correlations between adjacent learning days, with NE days 1-2 correlations subtracted (left), and the learning-associated change from NE days 1-2 to NE days 6-7 (right). Gray lines (right) indicate individual FOV × learning-environment observations (n = 20 from 10 FOVs × 2 environments; paired two-sided t-tests). **g.** Example spatial-field distributions along Track B during early and late learning for low- activity and high-activity neurons from one FOV. **h.** Landmark-aligned (top) and reward-aligned (bottom) spatial-field distributions for low- activity (left) and high-activity neurons (middle), alongside running speed (right), on NE day 1 and late learning (NE days 5–7). Black dots indicate significant differences between NE day 1 and late learning (paired two-sided t-tests with Benjamini-Hochberg correction). **i.** Schematic of the salient-feature score calculation. **j.** Landmark (left) and reward (right) salient-feature scores calculated from population spatial-field distributions of low-activity and high-activity neurons on NE day 1, days 2-4, and days 5-7 (n = 20: 10 FOV × 2 environments; paired two-sided *t*-tests). **k.** Changes in feature preference among neurons tracked from NE day 1 to late learning. Top left: transition matrices showing the percentage of low-activity neurons in each NE day 1 preference category (y axis) that transitioned to landmark-preferring, neutral, or reward-preferring on NE days 5-7 (x axis), so that the percentages across each row add up to 100% (n = 20 from 10 FOVs × 2 environments). Top right: similar to the left but for high-activity neurons. Bottom: difference between low- and high-activity groups using paired two-sided *t*-tests with Benjamini–Hochberg correction across the nine transitions. Blue indicates significantly greater transition probability in low-activity neurons, orange indicates significantly greater transition probability in high-activity neurons, and gray indicates no significant difference. **l.** Percentage of neurons remaining active across learning days grouped by activity and initial feature preference among neurons active on NE day 1 (n = 20 from 10 FOVs x 2 environments). Data are mean ± s.e.m. unless otherwise indicated. The dataset comprised 10 FOVs from 5 mice (2 FOVs per mouse), with each FOV contributing observations from two novel learning environments. Each FOV × learning environment was treated as one observation for the indicated FOV-level statistical comparisons (20 total observations).

However, additional features of low-activity neurons argued against this interpretation. Although they exhibited lower cross-run stability, quantified as the mean spatial activity correlation across all run pairs, this difference disappeared after activity matching (Fig. S3e). Nevertheless, low-activity neurons carried more spatial information (Fig. S3e) and showed greater learning-dependent improvements in both cross-run stability and spatial information. These greater learning-dependent improvements persisted in activity-matched data, indicating that spatial representations of low-activity neurons were not simply unstable (Figs. 2e and S3e). Both populations also remapped robustly upon novel environment exposure, demonstrating sensitivity to novel spatial contexts (Figs. S4a and b). As adjacent-session spatial activity correlations increased during learning, reflecting experience-dependent improvement in stability, low-activity neurons showed greater improvement, both before (Fig. 2f) and after activity matching (Fig. S3f). Together, these findings indicate that low-activity neurons encode spatial information and undergo greater learning-dependent refinement than high-activity neurons.

Importantly, the greater learning-dependent refinement of low-activity neurons was accompanied by a structured reorganization of their spatial field distribution, quantified as the percentage of neurons with fields at each track position. On the first day in a novel environment (Fig. 2g, Env. B), low-activity spatial fields preferentially concentrated near visual landmarks, whereas those of high-activity neurons were more broadly distributed (Fig. 2g, left). With learning, low-activity neurons showed a marked reduction in spatial fields near landmarks and an increase in the reward region, whereas high-activity neurons showed little reorganization (Fig. 2g, right). Feature-aligned population field distributions across environments B and C further revealed a progressive transition from landmark representation to pre-reward ramping in low-activity neurons during learning (Fig. 2h, left). In contrast, high-activity neurons showed relatively stable feature-associated representations, with reward activity closely tracking running speed (Fig. 2h, middle and right).

To quantify these dynamics, we calculated a salient-feature score, defined as the normalized difference between activity within and outside regions containing landmarks and reward. Higher scores indicate greater activity specificity near the corresponding features (Figs. 2i, 2h). In low-activity neurons, landmark scores were initially high and progressively decreased with learning, whereas reward scores increased. In contrast, high-activity neurons exhibited lower scores for both features, with little change across learning (Fig. 2j). These differences persisted after matching calcium events between low- and high-activity neurons (Fig. S4c) and were evident in both novel environments B and C when analyzed separately (Fig. S4d), indicating that the differential reorganization of feature-associated activity was not explained by differences in calcium-transient properties alone or by environmental identity. Moreover, when neurons were stratified more finely by relative activity, landmark scores in early learning and reward scores in later learning varied monotonically with relative activity (Fig. S4e), revealing a continuous relationship between feature-specific encoding and relative activity.

Finally, we examined how population-level reorganization emerged at the single- neuron level, distinguishing between two possibilities: individual neurons could shift their preferences from landmarks to reward (“within-cell reorganization”) and/or reward- preferring neurons could preferentially remain active (“selective stabilization”). Based on their relative landmark and reward scores on the first day in a novel environment, neurons were classified as landmark-preferring (higher landmark scores), reward-preferring (higher reward scores), or neutral (similar scores). Among low-activity neurons, a larger fraction of landmark-preferring and neutral neurons transitioned to the reward-preferring state, supporting within-cell reorganization. In contrast, high-activity neurons more often maintained their landmark-preferring or neutral identities, suggesting more stable feature tuning (Fig. 2k). Notably, the fraction of neurons that remained active as learning progressed did not differ according to their initial feature preference in either activity group (Fig. 2l), arguing against selective stabilization as the primary explanation for the population-level shift.

Together, these findings reveal complementary representational dynamics along the MEC activity continuum. High-activity neurons maintain relatively rich and stable spatial representations across learning, whereas low-activity neurons undergo greater experience-dependent representational refinement and reorganization. Importantly, this flexibility is structured: low-activity neurons shift from representing salient visual landmarks during early experience toward the learned reward location as learning progresses. Thus, lower-activity neurons provide a flexible component of the population representation that adaptively incorporates behaviorally relevant information during learning, while higher-activity neurons form a relatively stable component of the population representation.

### Pre-reward dopamine ramping links activity to reward representations

Because dopamine signals across multiple brain regions exhibit pre-reward ramping during goal-directed navigation, potentially providing a teaching signal for reward prediction^40–42^, we next examined whether dopamine dynamics were associated with the greater experience-dependent shift from landmark- to reward-specific representations in low- versus high-activity MEC neurons. We simultaneously imaged calcium activity and dopamine dynamics of excitatory neurons using a GRAB dopamine sensor rDA3m^43^ in GP5.3 mice (Figs. 3a, 3b). Mice learned a novel environment (NE) over seven days following familiar-environment (FE) training (Fig. 3c), with reward-predictive licking increasing in NE (Fig. 3d).

**Figure 3.**
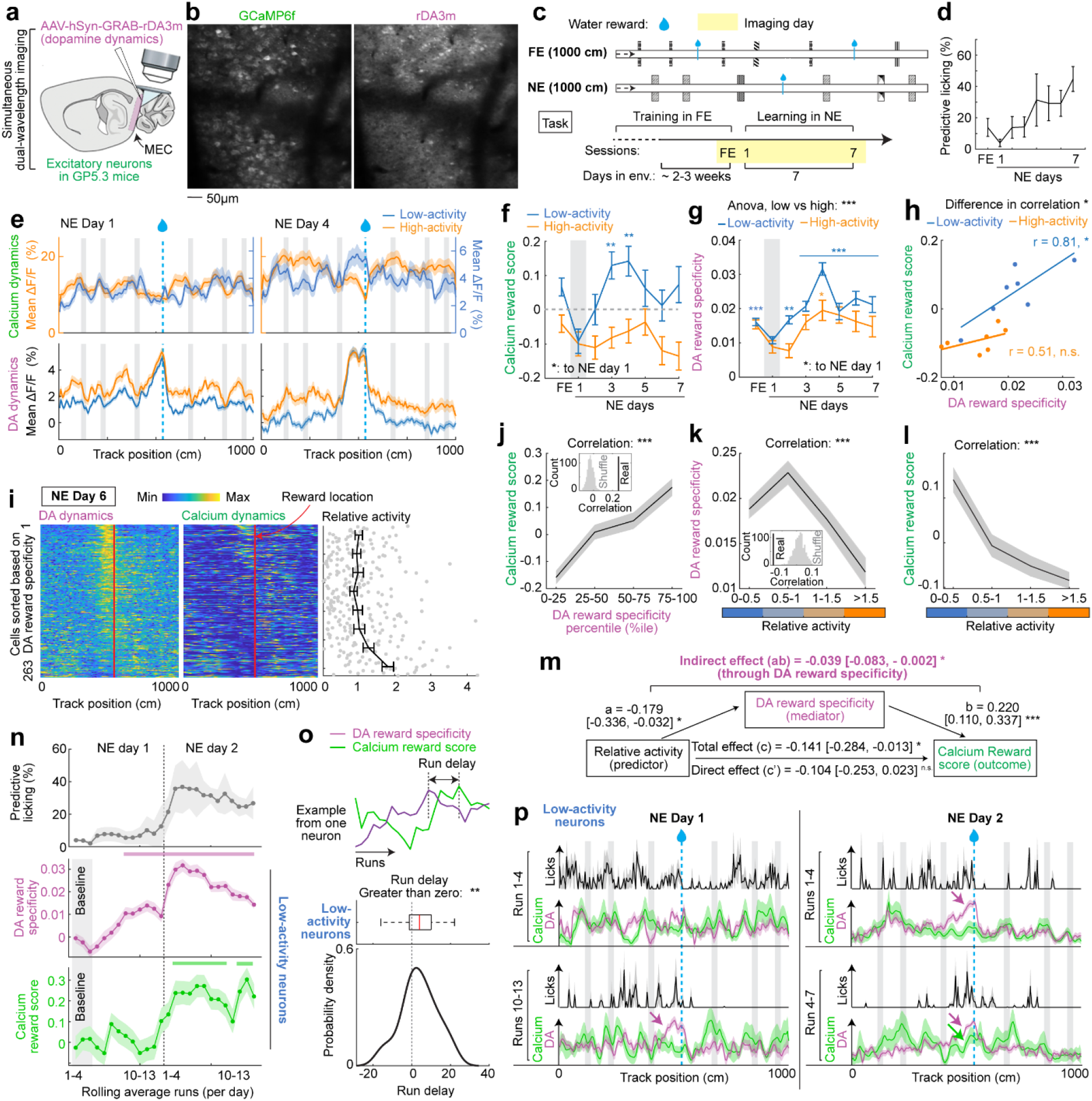
Dopamine pre-reward ramping links activity to reward representations. **a.** Schematic of simultaneous dual-wavelength imaging of MEC excitatory neurons (GCaMP6f) and dopamine dynamics (rDA3m) in GP5.3 mice. **b.** Representative FOV showing GCaMP6f⁺ neurons (green) and rDA3m expression (magenta). **c.** Track and task design. Mice were imaged on the last familiar-environment (FE) day and across 7 days in a novel environment (NE). **d.** Reward-predictive licking across FE and NE learning days (n = 3 mice). **e.** Spatially binned calcium activity of low- and high-activity neurons (top) and dopamine (DA) dynamics (bottom) on NE days 1 and 4. **f.** Calcium reward score by activity group. Gray indicates NE day 1, the reference for linear mixed-effects models (day fixed effect, FOV random effect), with Holm-Bonferroni correction. **g.** Similar to **f** but for DA reward specificity, defined as the difference between mean DA signals within 200 cm before reward delivery and those across the remainder of the track, excluding the 325 cm following reward delivery. Two-way ANOVA tested group differences. **h.** Correlation between day-averaged calcium reward score and DA reward specificity in low- and high-activity groups. Differences between correlation coefficients were tested against a null distribution generated by shuffling calcium reward scores within groups (1,000 iterations). **i.** Example of NE day 6 DA (left) and calcium (middle) dynamics along the track, and relative activity of individual neurons (right), sorted by DA reward specificity. **j.** Correlations between calcium reward score and DA reward specificity. Neuron-level measures were averaged across days when neurons were active. For visualization, calcium reward scores were binned by every 25^th^ percentile of DA reward specificity. Insets compare observed correlations with null distributions from 1,000 circular spatial shuffles of DA dynamics. **k.** Similar to **j** but for the correlation between DA reward specificity and relative activity. **l.** Similar to **j** but for the correlation between calcium reward score and relative activity. **m.** Mediation analysis showing a significant indirect effect of relative activity on calcium reward score through DA reward specificity, with a nonsignificant direct effect. Confidence intervals were obtained from 1,000 hierarchical bootstrap iterations, resampling mice and then neurons within mice. Variables were z-scored within mouse. **n.** Time course of reward-predictive licking (top), DA reward specificity (middle), and calcium reward score (bottom) on NE days 1-2, using low-activity neurons active on both days (n = 57 neurons from 6 FOVs in 3 mice). Values were averaged over a four-run rolling window. Significance was assessed against the baseline mean of rolling windows spanning runs 1-4, 2-5, and 3-6 using paired Student’s *t*-tests with Holm-Bonferroni correction; significant intervals are indicated by magenta (DA) and green (calcium) bars. **o.** Run delays in the emergence of calcium reward scores (example, top) and population- level delay distributions relative to zero (middle, bottom). A two-sided ttest was used to assess shifts relative to zero. **p.** Visualization demonstrating that DA pre-reward ramping (magenta arrow) emerged before calcium reward activity (green arrow). Note that DA pre-reward ramping started to emerge in later runs on NE day 1, but calcium reward activity only appeared in later runs on NE day 2. Data are mean ± s.e.m. unless otherwise indicated. *, *P* ≤ 0.05, **, **P** ≤ 0.01,***, ***P*** ≤ 0.001; n.s., *P* > 0.05. The dataset comprised 6 FOVs from 3 mice (2 FOVs per mouse). Relative activity, calcium reward score, and DA reward specificity were quantified at the single- neuron level.

Consistent with our previous observations, during novel environment learning, low- activity neurons gradually shifted from landmark- to reward-specific representations (Fig. 3e), reflected in decreasing landmark scores (Fig. S5a) and increasing reward scores (Fig. 3f). These experience-dependent changes were less pronounced in high-activity cells. In parallel, dopamine activity became increasingly reward-specific, characterized by pronounced pre-reward ramping followed by a rapid post-reward decline (Figs. 3e, g). Dopamine reward specificity, quantified as the difference in dopamine activity between the pre-reward region and regions away from reward, was greater for low- than for high- activity neurons. It also increased more rapidly in low-activity neurons during learning, becoming significantly higher than on NE day 1 by NE day 2. In contrast, high-activity neurons exhibited only a modest increase (Fig. 3g). Notably, the day-by-day increase in dopamine reward specificity closely tracked the increase in reward scores in low-, but not high-activity neurons (Fig. 3h). These observations led us to hypothesize that reward- specific dopamine signaling could help explain the relationship between relative neuronal activity and reward-specific representations.

To visualize this relationship, we ranked active neurons each day by their dopamine reward specificity (Fig. 3i, left). Neurons with higher dopamine reward specificity tended to exhibit greater calcium activity around reward, particularly in the pre- reward region, relative to other track locations (Fig. 3i, middle), despite having lower relative activity (Fig. 3i, right; Fig. S5b shows examples from other days). Consistent with these observations, dopamine reward specificity was positively correlated with calcium reward scores (Fig. 3j) and negatively correlated with relative activity, with both correlations differing significantly from shuffled controls (Fig. 3k). As expected from our previous findings (Fig. S4e), reward scores were also negatively correlated with relative activity (Fig. 3l).

We next performed mediation analysis to test whether dopamine reward specificity could statistically account for the relationship between relative neuronal activity and reward scores at the individual-neuron level: Relative activity was set as the predictor, dopamine reward specificity as the mediator, and calcium reward score as the outcome (Fig. 3m). The indirect effect through dopamine reward specificity was negative and significant (ab = −0.039, *p* < 0.05), whereas the direct effect of relative activity on calcium reward scores was not statistically significant after accounting for dopamine reward specificity. These results suggest that dopamine reward specificity statistically accounts for part of the association between relative neuronal activity and reward coding, consistent with the observation that neurons with lower relative activity exhibit higher dopamine reward specificity and stronger reward-related calcium encoding.

The differential dopamine dynamics associated with low- and high-activity neurons may be related to their anatomical organization (Fig. S2). Indeed, anatomical distance was positively associated with differences in relative activity, dopamine reward specificity, and calcium reward scores (Fig. S5c), suggesting am association between spatially organized dopamine dynamics, neuronal activity levels, and reward-related function.

Lastly, we asked whether the temporal dynamics of dopamine reward specificity were consistent with a potential teaching role in the emergence of reward-specific calcium activity in low-activity neurons. We focused on neurons that were active on the first two NE days (Fig. 3n). Dopamine reward specificity increased rapidly on NE day 1 and became significantly elevated above baseline (the initial three four-run rolling windows) during later runs (Fig. 3n, middle). In contrast, calcium reward scores did not become significantly elevated until NE day 2 (Fig. 3n, bottom), coinciding with improved reward- predictive licking on NE day 2 (Fig. 3n, top). At the individual-neuron level, increases in calcium reward scores also lagged behind the development of dopamine reward specificity (Fig. 3o). This temporal difference was further evident in run-by-run analysis, which showed that dopamine reward specificity emerged before detectable increases in calcium activity around reward (Fig. 3p).

Together, our results suggest that differences in dopamine reward specificity may contribute to the differential reward encoding of low- and high-activity MEC neurons. In low-activity neurons, the temporal precedence of dopamine reward specificity supports a model in which pre-reward dopamine ramping may serve as a teaching signal that contributes to the gradual reorganization of neuronal representations from landmark toward reward during learning.

### Interneuron coupling links activity to representational stability

While the above results implicated dopamine in the flexible representational reorganization of low-activity neurons, we next investigated potential mechanisms underlying the greater stability of high-activity neurons. We focused on inhibitory interneurons, which have been implicated in maintaining the stability of hippocampal representations^44,45^. In particular, we examined PV+ and SST+ interneurons, two interneuron populations that extensively interact with excitatory neurons in the MEC^46^.

To investigate interactions between interneurons and excitatory neurons, we simultaneously imaged their calcium dynamics by transgenically expressing GCaMP8m in PV+ or SST+ interneurons and virally expressing jRGECO1a^47^ in MEC layer 2 excitatory neurons (Figs. 4a, b). Water-restricted mice transitioned from a familiar environment (FE) to a novel environment (NE) (Fig. 4c), which they subsequently learned over seven days, as reflected by improved reward-predictive licking (Fig. 4d).

**Figure 4.**
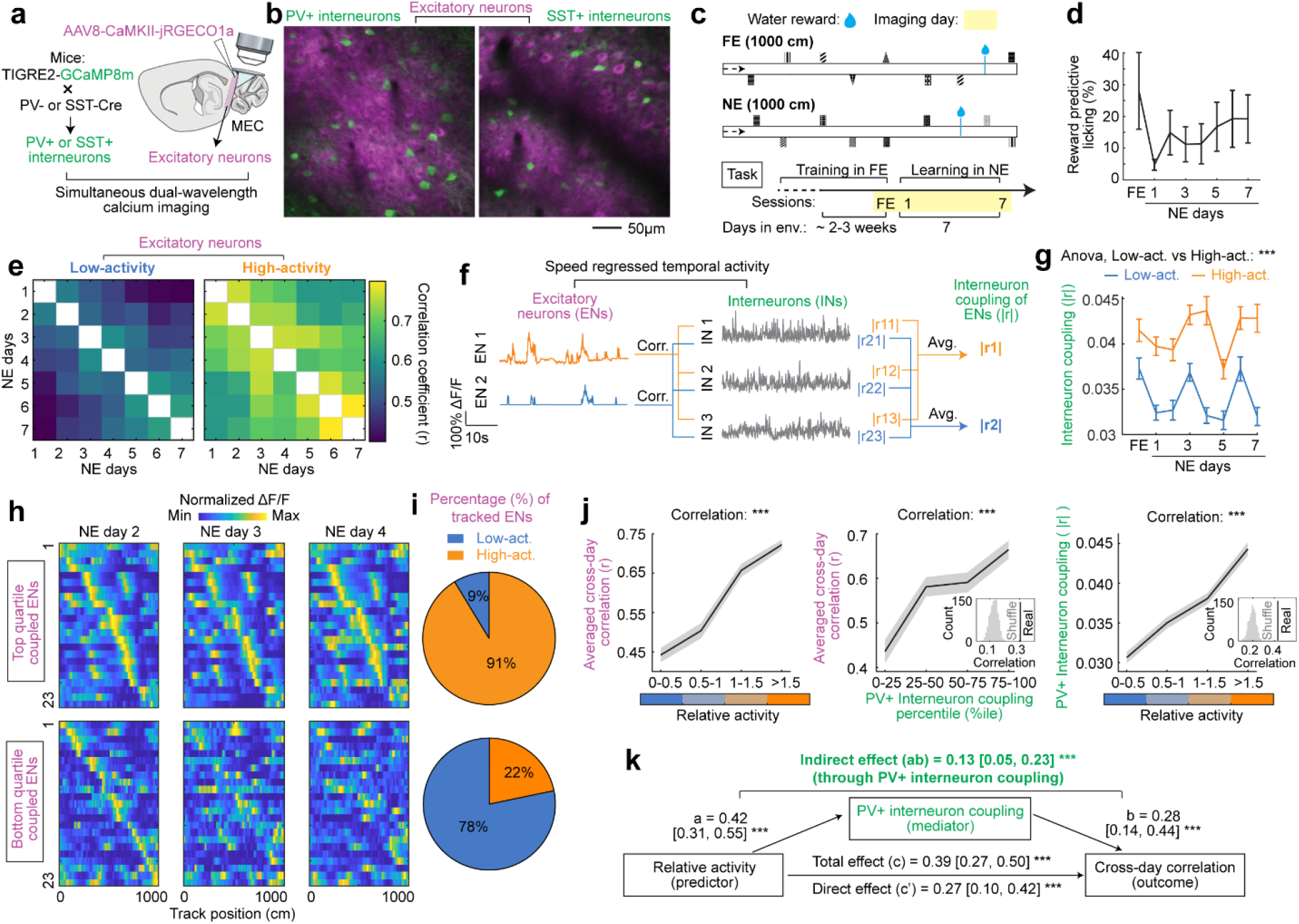
Interneuron coupling links activity to representational stability. **a.** Schematic of simultaneous dual-color calcium imaging of excitatory neurons and PV+ or SST+ interneurons in MEC. Excitatory neurons expressed jRGECO1a and PV+ or SST+ interneurons expressed GCaMP8m. **b.** Representative dual-color images showing excitatory neurons (magenta) and PV+ (left) or SST+ (right) interneurons (green). **c.** Experimental timeline. Following training and familiarization, mice were imaged once in the familiar environment (FE) and for 7 days in a novel environment (NE). **d.** Reward-predictive licking across FE and NE learning days (n = 4 mice). **e.** Mean pairwise spatial correlations across the 7 NE learning days for low-activity and high-activity excitatory neurons. Neurons were pooled across 4 FOVs from 4 mice. **f.** Schematic of excitatory–PV+ interneuron functional coupling analysis. Running speed was regressed from the temporal activity of both excitatory neurons and interneurons. Coupling for each excitatory neuron was calculated as the mean absolute Pearson correlation between its residual activity and that of all simultaneously imaged PV+ interneurons. **g.** Excitatory–PV+ interneuron coupling across learning days for low-activity and high- activity excitatory neurons. Neurons were pooled across 4 FOVs from 4 mice. An ANOVA tested the overall difference between low-activity and high-activity neurons. **h.** Spatial activity matrices of excitatory neurons in the top (top) and bottom (bottom) quartiles of PV+ interneuron coupling (averaged across all NE sessions) tracked through NE days 2, 3, and 4. Cells were sorted by peak location on NE2 and kept that ordering. Neurons were pooled across 4 FOVs from 4 mice. Top quartile group was downsampled to match the neuron count of the bottom quartile. **i.** Fraction of low-activity and high-activity excitatory neurons in the top (top) and bottom (bottom) quartiles of PV+ interneuron coupling for the displayed cells. **j.** Relationships among relative activity, PV+ interneuron coupling, and spatial stability. Left, spatial stability as a function of relative activity. Middle, spatial stability across quartiles of PV+ interneuron coupling. Right, PV+ interneuron coupling as a function of relative activity. Insets show corresponding circular permutation shuffle distributions. Significance was also assessed with a two-tailed Pearson correlation test. **k.** Mediation analysis testing whether PV+ interneuron coupling statistically mediates the relationship between relative activity and spatial stability. Total effect of relative activity on spatial stability, 0.39 [95% CI, 0.27, 0.50]; relative activity-to-coupling path, 0.42 [0.31, 0.55]; coupling-to-stability path controlling for relative activity, 0.28 [0.14, 0.44]; indirect effect through PV+ coupling, 0.13 [0.05, 0.23]; direct effect after accounting for PV+ coupling, 0.27 [0.10, 0.42]. Confidence intervals were obtained from 1,000 hierarchical bootstrap iterations, resampling mice and then neurons within mice. Data are mean ± s.e.m. unless otherwise indicated. The dataset comprised 4 FOVs from 4 mice (1 FOV per mouse). Relative activity, PV+ interneuron coupling, and spatial stability were quantified at the single-neuron level for analyses in **j**,**k**. Variables included in the mediation analysis were z-scored within mouse before analysis.

We first examined PV+ interneurons. Consistent with our previous observations, high-activity neurons exhibited greater spatial activity correlations across session pairs than low-activity neurons, indicating greater representational stability (Fig. 4e). To assess whether excitatory-interneuron interactions could explain this stability difference, we correlated the temporal activity of excitatory neurons and interneurons. Because the activity of both excitatory neurons and interneurons has been shown to be running-speed- modulated^48^, we regressed speed from their temporal activity before calculating correlations (Fig. 4f). For each excitatory neuron, functional coupling was defined as the mean absolute correlation with simultaneously imaged interneurons.

High-activity neurons exhibited stronger functional coupling with PV+ interneurons than low-activity neurons throughout learning (Fig. 4g). This difference was evident for both positive and negative correlations (Fig. S6a). Consistent with this relationship, excitatory neurons in the top quartile of PV+ coupling exhibited greater cross-day spatial stability than those in the bottom quartile (Fig. 4h). The top-coupling quartile also possessed a larger fraction of high-activity cells (Fig. 4i). Across the continuum of relative activity, PV+ coupling strength was positively correlated with both relative activity and cross-day spatial activity correlation (Fig. 4j). Circularly permuting excitatory activity in time substantially reduced these correlations (Fig. 4j, insets in middle and right panels), indicating that these associations depended on the temporal correspondence between excitatory and interneuron activity rather than excitatory activity magnitude alone.

These results suggested that PV+ coupling might statistically account for part of the association between relative activity and spatial stability. We therefore tested this relationship using mediation analysis, with relative activity as the predictor, PV+ coupling as the mediator, and spatial stability as the outcome. The indirect effect through PV+ coupling was positive and significant (ab = 0.13, 95% confidence interval [0.05, 0.23], p < 0.001) (Fig. 4k). Thus, PV+ functional coupling statistically accounted for part of the relationship between relative activity and spatial stability.

We next investigated whether similar relationships were present for SST+ interneurons. Like PV+ interneurons, SST+ functional coupling was greater for high- activity neurons than low-activity neurons (Fig. S6b), with a graded relationship across the continuum of relative activity (Fig. S6c, middle). However, the relationship between SST+ coupling and cross-day spatial activity stability did not reach statistical significance (Fig. S6c, right). Consistently, the indirect effect of SST+ coupling in the mediation analysis was small and nonsignificant (ab = 0.029, 95% CI [−0.033, 0.108], p > 0.05) (Fig. S6d). A direct comparison of the mediation effects of PV+ and SST+ functional coupling suggested a trend toward weaker effect of SST+ coupling, although the difference did not reach statistical significance (permutation test, P = 0.119; Fig. S6e). The small sample sizes (PV, n = 4 mice; SST, n = 5 mice) potentially limit the ability to resolve differences between the two interneuron populations.

Together, these findings reveal coordinated interactions between excitatory and inhibitory network activity during novel-environment learning. High-activity neurons exhibit stronger functional coupling with interneurons, and this stronger PV+ coupling is associated with greater spatial stability. These findings implicate excitatory-PV+ interneuron coupling as a potential network mechanism contributing to the stable representation of novel environments during learning.

### Initial reduction in inhibition facilitates learning-dependent remodeling of excitatory neural dynamics

While the above results revealed how flexible and stable representations might mechanistically evolve during learning, we next investigated how these dynamics are initiated upon novel environment exposure. While the magnitude of the initial increase in excitatory neural activity was closely associated with subsequent learning success (Fig. 1i), this increase was accompanied by decreased activity in both PV+ and SST+ interneurons (Fig. 5a, b). Reduced inhibition in other neural circuits can enhance excitatory neuron responsiveness and facilitate synaptic plasticity, providing a permissive state for experience-dependent circuit reorganization^49,50^. We therefore hypothesized that in the MEC, reduced inhibition upon novelty exposure facilitates the initial increase in excitatory neural activity, thereby promoting the subsequent development of flexible and stable excitatory representations that support successful spatial learning.

**Figure 5.**
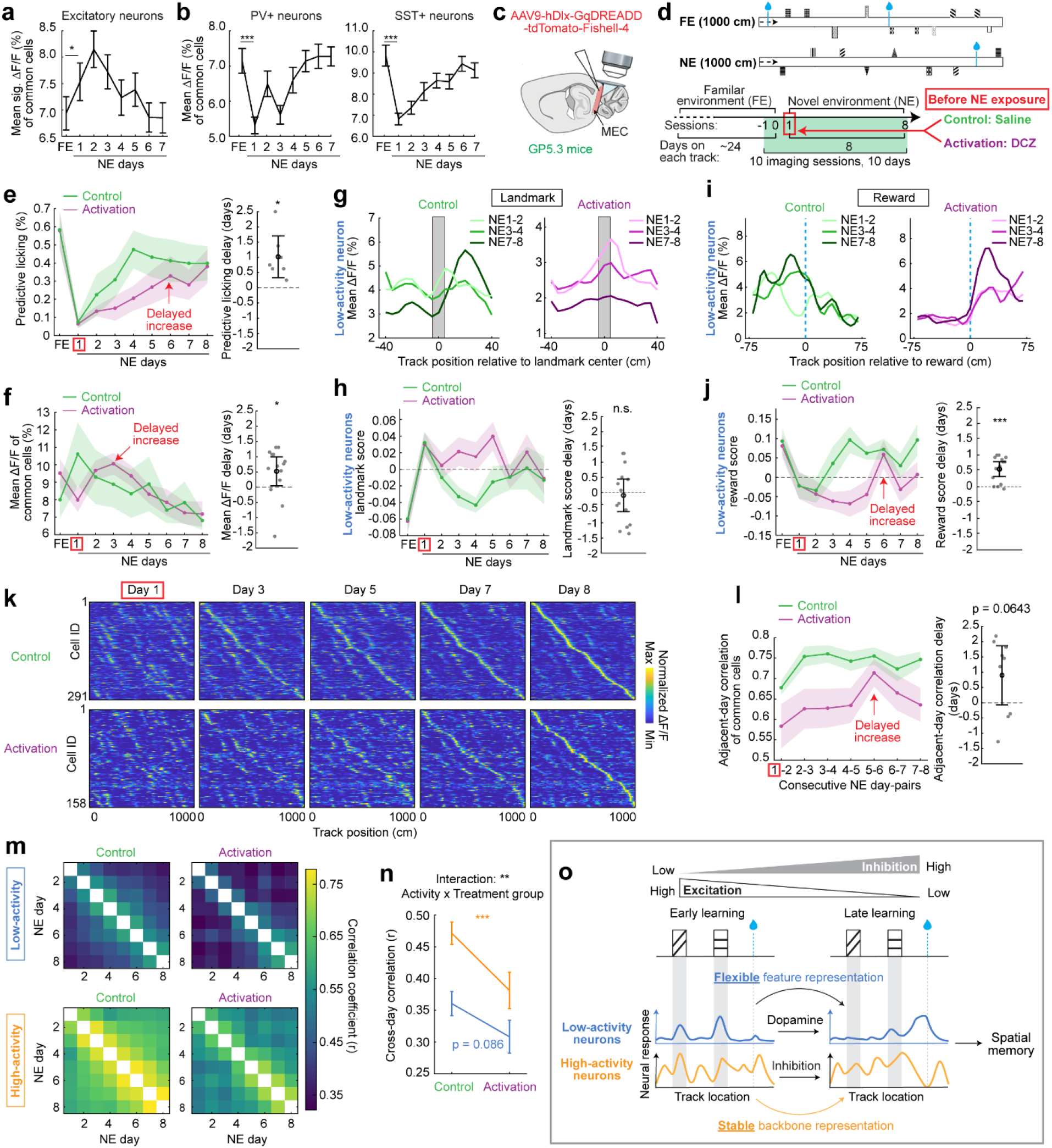
Initial reduction in inhibition facilitates learning-dependent remodeling of excitatory neural dynamics. **a.** Mean significant transient activity of excitatory neurons across familiar-environment (FE) and novel-environment (NE) sessions. Neurons were pooled across 9 fields of view (FOVs) from 9 mice. **b.** Calcium activity (ΔF/F) of PV+ and SST+ interneurons across FE and NE sessions. PV+ and SST+ interneurons were pooled across 4 FOVs from 4 mice and 5 FOVs from 5 mice, respectively. **c.** Schematic illustrating viral expression of hM3Dq(Gq)-dTomato in MEC inhibitory interneurons of GP5.3 mice for chemogenetic activation of interneurons before novel- environment exposure. **d.** Experimental paradigm: Mice were trained in a familiar environment before being exposed to a novel environment, which they subsequently learned over eight days. Mice received DCZ or saline 10 minutes before initial NE exposure. **e.** Reward-predictive licking across learning in saline-treated control and DCZ-treated activation mice (left), and estimated temporal delays in the activation group obtained from the max cross-correlation with control trajectories (right). n = 8 control mice and 7 activation mice. **f.** Mean excitatory neuronal activity for neurons imaged across all FE and NE sessions in control and activation groups (left), and estimated temporal delays in the activation group obtained from the max cross-correlation with control trajectories (right). n = 14 FOVs from 7 control mice and 14 FOVs from 7 activation mice. **g.** Example FOV spatial activity of low-activity neurons aligned to landmark locations across learning in control and activation groups. **h.** Landmark scores of low-activity neurons across learning in control and activation groups. Scores were calculated per neuron and aggregated within each FOV. Right, estimated temporal delays in the activation group relative to controls. **i.** Same as **g**, but for spatial activity aligned to the reward location. **j.** Same as **h**, but for reward scores. **k.** Example spatial activity heatmaps of longitudinally tracked excitatory neurons across learning in control (top) and activation (bottom) groups. Neurons were sorted by their peak spatial activity on NE day 8. **l.** Adjacent-day spatial activity correlations of neurons active across all sessions in control and activation groups (left), and estimated temporal delays in the activation group relative to controls (right). **m.** Cross-day spatial activity correlation matrices across eight NE learning days for low- activity and high-activity neurons in control and activation groups. **n.** Quantification of cross-day spatial stability in low-activity and high-activity neurons across control and activation groups. Group differences and their interaction were assessed with a two-way ANOVA. **o.** Proposed model illustrating how the initial reduction in interneuron activity during novel- environment exposure facilitates the experience-dependent reorganization and stabilization of excitatory spatial representations supporting spatial learning. Data are presented as mean ± s.e.m. unless otherwise indicated.

To test this hypothesis, we asked whether activating MEC interneurons on the first day of novel-environment exposure, thereby counteracting the natural interneuron activity reduction, would disrupt the subsequent development of excitatory neural representations. We virally expressed AAV-hDlx-hM3Dq(Gq)-dTomato in GP5.3 mice to target MEC inhibitory interneurons, mostly including PV+ and SST+ cells^51^ (Fig. 5c). After training in a familiar environment (FE), mice were assigned to activation or control groups before being exposed to the novel environment (NE). Ten minutes before NE exposure, mice in the activation group received deschloroclozapine (DCZ)^52^, whereas control mice received saline (Fig. 5d). We longitudinally tracked GCaMP6f activity from the final two days in FE through the first eight days of NE learning. In a separate cohort, we confirmed that DCZ increased PV+ and SST+ interneuron activity during navigation (Fig. S7).

We first examined whether this transient interneuron activation affected learning. Compared with the controls, mice in the activation group showed a later increase in reward-predictive licking, although late-learning performance did not significantly differ between groups (Fig. 5e, left). Cross-correlation of the groups’ learning trajectories further supported a temporal shift in the activation group (Fig. 5e, right), indicating that transient interneuron activation delayed, rather than prevented, spatial learning.

To characterize the excitatory neuronal dynamics associated with this delayed learning, we examined changes in neural activity levels across days. In the control group, excitatory activity increased during initial NE exposure and subsequently declined with experience, as expected. In contrast, the initial activity increase on NE day 1 was suppressed in the activation group (Fig. 5f, left), and this suppression was observed in both low- and high-activity neurons (Fig. S8a). Notably, elevated activity emerged on subsequent learning days and persisted for several days, suggesting a delayed activity response. Indeed, cross-correlation of the excitatory activity trajectories showed a temporal shift in the activation group (Fig. 5f, right). Thus, transient interneuron activation during initial novelty exposure delayed the normal progression of excitatory neural dynamics across learning.

We further examined how interneuron activation affected the spatial activity of excitatory neurons, focusing first on remapping upon novel-environment exposure. In the control group, spatial activity patterns were poorly correlated between the final FE day and the first NE day, consistent with robust remapping. In contrast, the correlation distributions in the activation group were positively shifted, indicating reduced remapping (Fig. S8b). Although the shifts did not differ significantly across activation condition between high- and low-activity neurons (Fig. S8c), a significant shift was detected in high- activity neurons (Fig. S8b, bottom), suggesting that they were more susceptible to the activation effect, consistent with their stronger coupling to interneurons (Fig. 4). Together, these results indicate that interneuron activation was associated with reduced high- activity neuron remapping upon novel environment exposure.

We next asked whether interneuron activation altered the flexible coding of salient features by low-activity neurons. In controls, these neurons initially displayed landmark coding that decreased with experience (Fig. 5g, h, left), while reward coding increased significantly up to NE day 4 (Fig. 5i, j, left). In the activation group, the initial landmark coding was comparable to that in controls but remained elevated through approximately NE day 5 (Fig. 5g, right, and 5h), accompanied by a weaker and delayed increase in reward coding (Fig. 5i, right and 5j). These findings suggest that transient interneuron activation delayed the experience-dependent reorganization of low-activity neural representations from landmarks toward reward.

Finally, we examined whether interneuron activation affected the stabilization of spatial representations. In controls, spatial representations stabilized rapidly across adjacent days (Fig. 5k, top and 5l). In contrast, the activation group showed a trend toward delayed stabilization (Fig. 5k, bottom and 5l). Across learning, the activation group also exhibited reduced representational stability, with a more pronounced reduction in high- activity neurons (Fig. 5m and n). Notably, within this group, lower stability was associated with less remapping on NE day 1 (Fig. S8d), with a stronger relationship in high- than in low-activity neurons (Fig. S8e). This relationship suggests that proper remapping upon novelty exposure may facilitate subsequent stabilization during learning. The significantly reduced remapping of high-activity neurons may therefore contribute to their greater reduction in stability. Together, these results show that transient interneuron activation upon novelty was associated with delayed stabilization and reduced overall stability of spatial representations, particularly in high-activity neurons.

Together, these findings link the initial reduction in inhibition upon novel- environment exposure to the subsequent progression of excitatory MEC neural dynamics and spatial learning. Transiently preventing this reduction through interneuron activation delayed learning and was associated with alterations in several features of excitatory neural activity, including suppression of the initial activity increase, reduced remapping, delayed experience-dependent changes in excitatory neural activity, delayed reorganization of low-activity neural representations from landmarks to reward, and delayed and reduced stabilization of spatial representations. These findings suggest that the initial reduction in inhibition upon novelty exposure facilitates the experience- dependent formation of flexible and stable spatial representations by excitatory neurons, a process that parallels the progression of spatial learning and enables the formation of effective spatial memory (Fig. 5o).

## Discussion

Successful memory formation requires neural circuits to incorporate novel information while stabilizing acquired representations. Here, we identify coordinated neuronal mechanisms in the MEC that may support these seemingly opposing demands (Fig. 5o). MEC neurons span a skewed continuum of activity that is preserved across environments and learning stages. Low-activity neurons are preferentially recruited by novelty, exhibit more intermittent activation across learning, and show greater learning- dependent refinement, shifting from landmark- to reward-related representations. By contrast, high-activity neurons remain more persistently active and carry richer, more stable spatial representations. These complementary encoding properties are potentially mediated by dopaminergic signaling and differential functional coupling with interneurons. Reward-specific dopamine ramping is more pronounced in low-activity neurons and precedes the emergence of their reward-related coding, whereas stronger interneuron coupling with high-activity neurons is associated with greater representational stability. The emergence of these complementary properties appears to be gated by an initial reduction in inhibition, as transiently preventing this reduction during initial exposure to a novel environment delayed novelty-associated excitatory responses, the development of flexible and stable representations, and behavioral learning. Together, these findings identify activity-dependent neuronal organization and circuit modulation as candidate mechanisms through which the MEC balances representational flexibility with stability to support spatial learning.

Skewed distributions of neuronal activity have been reported across multiple brain regions^37,53–55^ and species^37,56,57^, and are preserved across environments and brain states^37,53,58,59^, as described in our current study, suggesting that this distribution is a general feature of neural networks^57^. Along this activity continuum, highly active neurons have been shown to carry rich and stable information^37,60^, whereas low-activity neurons exhibit greater plasticity^38^ and have been proposed to provide the flexibility needed for dynamic neural encoding^57^. However, how this flexibility contributes to information encoding during spatial learning remains unclear. In the hippocampus, low-activity neurons exhibit robust representational drift even in familiar environments^13^, further raising the question of whether their flexibility reflects selective information updating or simply nonspecific representational instability. Here, we show that low-activity MEC neurons exhibit not only greater experience-dependent refinement but also structured representational remodeling that closely tracks changes in behaviorally relevant information during learning. Upon initial exposure to a novel environment, these neurons rapidly developed landmark-related representations, preferentially encoding salient environmental features. As learning progressed and the reward location acquired behavioral significance, their representations shifted toward reward-related coding. Thus, the flexibility of low-activity neurons reflects selective, experience-dependent updating of behaviorally relevant information rather than simply degraded spatial coding. These findings suggest that low-activity neurons preferentially support context- and experience- dependent representations, whereas high-activity neurons provide a more stable, information-rich representational scaffold of the environment. Although these properties vary along a continuum of neuronal activity, their differences are not fully explained by activity level alone, suggesting that additional functional properties contribute to representational diversity. Moreover, the fewer spatial fields and higher salient-feature scores of low-activity neurons suggest a sparse, selective coding regime that prioritizes salient environmental features. Sparse coding has been observed in the hippocampus^53,61^ and sensory systems^62^ and has been proposed to increase memory capacity by limiting overlap among stored representations, thereby reducing interference between experiences^63,64^. The selective coding of low-activity neurons may therefore favor the formation of context-specific representations. Together, the complementary representational dynamics of low- and high-activity neurons may enable the MEC to incorporate behaviorally relevant new information while maintaining coherent, environment-specific spatial representations during learning.

Beyond their functional differences, low- and high-activity neurons also differ in their anatomical organization and association with putative molecular cell types. Differences in relative activity increased with inter-somatic distance, indicating that neurons with similar activity levels tended to be spatially clustered. This organization contrasts with the salt-and-pepper arrangement of neurons with different activity levels reported in the hippocampus^53^. Such anatomical clustering may reflect, or contribute to, differences in local connectivity within excitatory neurons, their interactions with interneurons, and access to long-range inputs, potentially shaping the distinct representational dynamics of the two populations. Notably, the anatomical organization of neurons according to relative activity was accompanied by a similar spatial organization of dopamine reward specificity. This correspondence raises the possibility that spatial variation in dopamine signaling contributes to the activity and functional organization of MEC neurons. Given evidence that dopamine modulates neuronal excitability and other neuronal functions in the entorhinal cortex^65–69^, local differences in dopamine signaling could contribute to the activity and coding properties of these populations, although this possibility remains to be tested mechanistically. Furthermore, low- and high-activity neurons also differ in their association with putative MEC layer 2 cell types. Although putative stellate cells predominate in both populations, low-activity neurons contain a higher proportion of putative pyramidal cells, which differ from stellate cells in their intrinsic electrophysiological properties and connectivity within and beyond the MEC^46,70^. These differences in cell-type composition may contribute to the functional diversity observed along the activity continuum. Collectively, these findings suggest that the functional organization of low- and high-activity neurons may reflect the combined influence of anatomical organization, circuit connectivity, neuromodulatory inputs, and intrinsic cellular properties. This interpretation is consistent with broader evidence linking neuronal activity to synaptic strength, cortical connectivity, and axonal properties^57^.

Our findings further suggest that MEC interneurons regulate excitatory representational dynamics at both the population and individual-neuron levels. At the population level, interneurons may temporally gate excitatory plasticity during initial exposure to a novel environment. While excitatory neuronal activity initially increases and then gradually declines during learning, interneuron activity exhibits an opposing trajectory, decreasing upon novelty exposure before progressively recovering. This transient reduction in interneuron activity may create a permissive window for experience- dependent remodeling of excitatory representations^49,50^. Consistent with this hypothesis, preventing the novelty-associated reduction in interneuron activity suppressed excitatory neuronal activity and impaired remapping, suggesting that this initial inhibitory response contributes to appropriate novelty-dependent changes in neural dynamics. Moreover, interneuron activation delayed the learning-dependent evolution of excitatory activity levels, the transition from landmark- to reward-related representations in low-activity neurons, representational stabilization, and spatial learning. The coordinated delays across neural and behavioral measures suggest that the initial reduction in interneuron activity is important for the timely emergence of flexible and stable representations that support spatial learning. Novelty-associated reductions in inhibition have also been implicated in facilitating synaptic plasticity and the formation of task-relevant representations in other brain regions, including the hippocampus^44,71–73^, motor cortex^74^, and sensory cortices^75–77^. While our findings suggest that both PV+ and SST+ interneurons contribute to the novelty-associated reduction in MEC interneuron activity, the underlying mechanisms remain unclear. Potential contributors include neuromodulatory inputs, particularly cholinergic signaling^78^, and disinhibition mediated by VIP (vasoactive intestinal polypeptide)+ interneurons, which suppress other inhibitory populations and thereby reduce inhibitory drive onto excitatory neurons^79,80^.

At the individual-neuron level, interneurons may differentially regulate the stability of excitatory neurons during learning according to their relative activity levels. Weaker interneuron coupling with low-activity neurons may permit greater representational flexibility, consistent with their more pronounced improvement in spatial representation consistency and stronger transition from landmark- to reward-related coding. Conversely, stronger interneuron coupling with high-activity neurons may help constrain representational changes and preserve their information-rich representations. Consistent with this possibility, enhancing interneuron activity during novelty exposure disrupted remapping more robustly in high-activity neurons, suggesting that stronger inhibition may constrain their ability to update representations in response to novelty. This disruption may, in turn, contribute to their greater impairment in subsequent representational stabilization, as evidenced by the observed correlation between the extent of remapping on NE day 1 and representational stability during learning. In line with our findings, differential innervation of interneurons to excitatory neurons has been proposed to be associated with variations in their firing rate^57^. Together, these findings suggest that activity-dependent differences in interneuron coupling may help balance representational flexibility and stability in the MEC.

Lastly, while inhibitory signaling may regulate the timing and stability of representational remodeling, our findings suggest that dopamine may contribute to the flexible updating of MEC representations by biasing this remodeling toward behaviorally relevant information, particularly reward. The entorhinal cortex receives dopamine inputs from the ventral tegmental area (VTA) in the midbrain^81^ and may also receive dopamine released alongside noradrenaline from the locus coeruleus (LC)^82–84^. Dopamine ramping during goal-directed navigation has been observed in midbrain and striatal circuits and has been proposed to provide a teaching signal for reward-guided learning^41,42,85^. Recent work also identified pre-reward dopamine ramping in the hippocampus during spatial goal-directed behavior^40^. Consistent with these findings, we observed pre-reward ramping in extracellular dopamine signals in the MEC, with greater reward specificity associated with low-activity neurons and a close association with their progressive shift from landmark- to reward-related coding. Notably, dopamine ramping emerged before reward-related calcium coding in low-activity neurons, raising the possibility that dopamine helps guide the incorporation of reward-related information into evolving spatial representations, although a causal role remains to be established. In high-activity neurons, however, lower reward-ramping specificity of dopamine appeared to reflect higher background signals in non-reward regions rather than a weaker reward-related increase per se. This heterogeneity in dopamine dynamics associated with neuronal activity levels could arise through several, non-mutually exclusive mechanisms. One possibility is that low- and high-activity neurons receive different dopaminergic inputs. VTA- and LC-derived projections to the hippocampus exhibit distinct dynamics during goal-directed navigation^86^, raising the possibility that these inputs may also contribute differently to dopamine dynamics in MEC neurons. Alternatively, the observed differences may reflect local variation in extracellular dopamine dynamics, arising from differences in dopamine release, diffusion, or clearance. The anatomical segregation of low- and high- activity neurons potentially facilitates these differences under either scenario. However, because our sensor measures extracellular dopamine^43^, our data cannot distinguish among these possibilities. Further studies are therefore needed to resolve the underlying mechanisms. Nevertheless, our findings suggest complementary roles for inhibitory and dopaminergic signaling in learning-dependent plasticity, whereby inhibitory circuits gate when representational remodeling occurs and dopaminergic signaling influences what behaviorally relevant information is incorporated into flexible representations. In different tasks, other neuromodulatory or circuit mechanisms may support the flexible incorporation of behaviorally relevant information^87,88^, allowing cognitive maps to adapt to different task demands.

In summary, our study provides a framework for how coordinated excitatory, inhibitory, and neuromodulatory mechanisms may support the balance between representational flexibility and stability during spatial memory formation. These findings suggest a potential general principle of adaptive learning in which neural circuits incorporate behaviorally relevant information while preserving stable representations.

## Acknowledgement

We thank all members in the Gu laboratory for supporting the work, Dr. Ila Fiete, Dr. Lorna Role, Dr. David Talmage, and Dr. Mark Wagner for constructive suggestions on the project, and Dr. Chris McBain for supporting Dr. June Hoan Kim during the execution of this work. This research was supported by the Intramural Research Program of the National Institutes of Health (NIH) to Yi Gu (ZIA NS009415) and to Chris McBain (Z1A- HD001205-32). The contributions of the NIH authors were made as part of their official duties as NIH federal employees, are in compliance with agency policy requirements, and are considered Works of the United States Government. However, the findings and conclusions presented in this paper are those of the authors and do not necessarily reflect the views of the NIH or the U.S. Department of Health and Human Services.

## Author contribution statement

Conceptualization: DY, TM, YG

Methodology: DY, YM, LC, TM, YG

Investigation: DY, YM, LC, GW, TM, KZ, JHK, FS, KC, EB, YG

Visualization: DY, LC, YG

Funding acquisition: YG

Project administration: YG

Supervision: YG

Writing - original draft: DY, YG

Writing - review & editing: DY, YG

## Competing interests

The authors declare no competing interests.

## Supplementary Materials

### Materials and Methods

#### Animals

All animal procedures were performed in accordance with animal protocol 1524 approved by the Institutional Animal Care and Use Committee (IACUC) at NIH/NINDS. For the main excitatory neuron calcium imaging experiments, we used 3 male and 2 female GP5.3 mice^35^ (C57BL/6J-Tg(Thy1-GCaMP6f) GP5.3Dkim/J, JAX stock #028280) aged 4-5 months at the time imaging began. For simultaneous imaging of dopamine and excitatory neuron dynamics, we used 3 male GP5.3 mice injected with AAV9-hSyn- rDA3m. Mice were approximately 3-4 months old at the beginning of imaging. For simultaneous imaging of PV+ or SST+ interneurons and excitatory neurons, we used 2 male and 2 female TIGRE2+/−;PV-cre+/− mice or 2 male and 3 female TIGRE2+/−;SST- cre+/− mice, respectively, injected with AAV8.CamKIIa.NES.jRGECO1a.WPRE.SV40. Mice were approximately 4-5 months old at the beginning of imaging. For chemogenetic experiments, GP5.3 mice injected with AAV9-hDlx-GqDREADD-dTomato-Fishell-4 were used. Following the exclusion of one mouse (see Chemogenetic analyses), The deschloroclozapine (DCZ) group consisted of 4 male and 3 female mice, and the saline control group consisted of 4 male and 4 female mice. Mice were approximately 3-7 months old at the beginning of imaging. Mice were maintained on a reverse 12-h light/dark cycle, with all experiments performed during the dark period. Animals were housed at a temperature of 70-74 °F and 40-65% humidity. Analyses were not separated by sex because there was no a priori hypothesis that the experimental effects would differ by sex.

### Surgery

#### Microprism construction

Microprism construction was performed as described previously^89^. A cannula (MicroGroup, 304H11XX) was attached to a circular cover glass (3 mm, Warner Instruments, 64-0720). A right-angle microprism with an aluminum-coated hypotenuse (1.5 mm, OptoSigma, RBP3-1.5-8-550) was attached to the opposite side of the cover glass. All attachments were performed using UV-curing optical adhesive (ThorLabs, NOA81).

#### Microprism implantation surgery

Microprism implantation was performed as described previously^89^. Mice were anesthetized using a tabletop laboratory animal anesthesia system (induction: 3% isoflurane, 1 L/min oxygen; maintenance: 0.5–1.5% isoflurane, 0.7 L/min oxygen; VetEquip, 901806), and surgery was performed using a stereotaxic alignment system (Kopf Instruments, 1900). A homeothermic pad and monitoring system (Harvard Apparatus, 50-7220F) was used to maintain body temperature at 37 °C. After anesthesia induction, dexamethasone (2 mg/kg, VetOne, 13985-533-03) and saline (500 µL, 0.9% NaCl, McKesson, 0409-4888-50) were administered by intraperitoneal (IP) injection, and slow-release buprenorphine (1 mg/kg, ZooPharm, Buprenorphine SR-LAB) was administered subcutaneously. Enroflox 100 (10 mg/mL, VetOne, 13985-948-10) was used as an antimicrobial wash after the skull was exposed and again immediately prior to sealing the skull. All implants were placed in the left hemisphere, consistent with previous observations of more favorable vasculature on the left side^89^. A 3-mm craniotomy was centered 3.4 mm lateral to the midline and 0.75 mm posterior to the center of the transverse sinus (approximately 5.4 mm posterior to bregma). Mannitol (3 g/kg, Millipore Sigma, 63559) was administered intraperitoneally prior to a durotomy performed over the cerebellum. The microprism was inserted into the transverse sinus and sealed to the skull with Vetbond (3M, 1469SB). A headplate was then mounted on the skull opposite the craniotomy. Finally, the prism and headplate were secured to the skull with Metabond (Parkell).

### Virus injections

Mice underwent stereotaxic viral injections into the medial entorhinal cortex (MEC) with the head tilted upward by 18°. Injections were performed at two sites: 0.77 mm anterior to the transverse sinus, 3.00 mm lateral to bregma, and 1.79 mm below the brain surface; and 0.60 mm anterior to the transverse sinus, 3.36 mm lateral to bregma, and 1.42 mm below the brain surface. All viruses were pressure-injected through a glass micropipette at 100 nL/min. For interneuron imaging, mice received unilateral injections into the left MEC of AAV8-CamKIIa-NES-jRGECO1a-WPRE-SV40 (NEI Virus Packaging Core; 1 × 10¹³ vg/mL, diluted fourfold in mannitol), with 200 nL injected per site. For chemogenetic experiments, mice received bilateral injections of AAV9-hDlx-GqDREADD- dTomato-Fishell-4 (Addgene; 3.4 × 10¹³ vg/mL, diluted tenfold in mannitol), with 100 nL injected per site. For dopamine imaging, mice received unilateral injections into the left MEC of AAV9-hSyn-rDA3m (2 × 10¹² vg/mL, diluted twofold in mannitol), with either 100 or 200 nL injected per site.

### Virtual reality

For all behavioral experiments, a customized virtual reality (VR) setup was used, similar to that described previously^18^. Mice were head-fixed using the mounted headplate above an air-supported polystyrene ball (8” diameter, Smoothfoam). The ball rotated on an axle, allowing only forward and backward rotation. The virtual environment was projected onto a dome screen that covered approximately 270° of the visual field. An optical flow sensor (Paialu, paiModule_10210) with infrared LEDs (DigiKey, 365-1056- ND) was used to measure rotation of the ball and control movement through the virtual environment. The optical flow sensor sent its output to an Arduino board (Newark, A000062), which transmitted the movement signal to the computer controlling the VR. Water rewards of approximately 4 µL were delivered at fixed locations using a lick tube and solenoid. Licking was detected using a lick sensor connected to the lick tube and headplate holder. Licking the tube closed a circuit through the mouse, headplate, and headplate holder. The solenoid and lick sensor were controlled using a Multifunction I/O DAQ (National Instruments, PCI-6229). Virtual environments were generated and projected using ViRMEn software^90^ (Princeton, version 2016-02-12) in MATLAB R2015aSP1. ViRMEn environments were updated at 60 Hz and the DAQ input/output rate was 1 kHz. Imaging and behavioral data were synchronized by recording voltage signals corresponding to behavioral parameters from the VR system using the DAQ. The synchronization voltage signal was updated at 20 kHz, and behavioral outputs were matched to the imaging data for subsequent analysis. Projected light was passed through a Wratten filter (Kodak, 53-700) to reduce contamination of the imaging signal. Virtual environments consisted of 1D linear tracks with patterned walls and visual landmarks at fixed locations. At the end of each track, mice were immediately teleported back to the beginning.

### Behavior

#### Training in VR

Mice were allowed to recover for 5 days following microprism implantation and were then placed on water restriction, receiving 1 mL of water per day. After approximately 3 days of water restriction, mice began daily training in VR. Mice were first trained on a 1-m track to encourage running and were then trained on a 10-m track containing two reward locations until they showed consistent running (>40 trials per hour) and stable anticipation of reward locations for 3 consecutive days, defined as less than a 5% change in predictive licking.

For the main excitatory-neuron dataset, mice were imaged once in the familiar environment (Track A, 1000 cm) and then switched to a novel environment (Track B, 875 cm), which they navigated for 7 days. Following imaging on day 7 in Track B, mice were exposed to a second novel environment (Track C, 875 cm), which they subsequently navigated for 7 days. Imaging sessions lasted approximately 45 min and contained at least 12 laps. The same novel-environment paradigm was repeated in the other experiments, although the tracks and number of novel-environment imaging days differed.

#### Reward-predictive licking

Predictive licking was quantified as the percentage of licks occurring within a defined distance before the reward location (“pre-reward distance”), relative to licks at all other track positions, excluding those occurring within a defined distance after reward delivery (“post-reward distance”) to eliminate reward-consumptive licking^18^. These distances varied across datasets according to track design and mouse behavior: 25 cm and 15 cm, respectively, for Fig. 1; 20 cm and 30 cm for Fig. 3; 50 cm and 40 cm for Fig. 4; and 20 cm and 50 cm for Fig. 5. For Fig. 4, analyses were restricted to the first 10 runs because this dataset typically included more runs (>30) than the others, and many animals showed reduced attention to reward delivery during later runs.

#### Running speed

Mean running speed was calculated across the entire imaging session, including periods of immobility. Running speed before and after exposure to each novel environment was compared across mice using paired two-sided t-tests.

### Two-photon imaging

In general, imaging was performed using a microscope (Bruker) integrated with the VR setup described above. Imaging was performed using a 16× water-immersion objective (Nikon, MRP07220), with ultrasound transmission gel (Sonigel, Mettler Electronics) used as the immersion medium. Laser scanning was performed using a resonant-galvo scanner (Cambridge Technology, CRS8K). Before the first imaging session, the anterior-posterior and medial-lateral angles of the prism were measured relative to the head-fixed position. Before each subsequent session, the headplate holder and objective angle were adjusted to align the objective with the prism surface. The objective and imaging window were enclosed with black tubing to reduce contamination from projected light. Images were acquired using Prairie View 5.7 (Bruker) and converted using the Bruker Image-Block Ripping Utility.

#### Single-channel GCaMP6f imaging in GP5.3 mice: multi-environment experiments

GCaMP6f imaging was excited using Chameleon Discovery NX laser (Coherent) tuned to 920 nm, and the fluorescence was isolated using a 525/25-nm emission filter and detected using GaAsP photomultiplier tubes (Hamamatsu, H10770PB). Two planes separated by 30 µm within layer 2 were imaged using a Z-piezo. Each plane was acquired at 7.4 Hz with a resolution of 512 × 512 pixels (1.116 µm/pixel, full FOV 571.4 µm × 571.4 µm). Average laser power at the front of the objective was approximately 70-115 mW. Imaging and behavioral recordings were synchronized as described above.

#### Single-channel GCaMP6f imaging in GP5.3 mice: chemogenetic experiments

The imaging was conducted on two rigs, where GCaMP6f imaging was excited using either a Chameleon Discovery NX laser (Coherent) tuned to 920 nm (rig 1), or a 920-nm fixed-wavelength laser (Axon, Coherent) (rig 2), and the fluorescence was isolated using a 525/25-nm emission filter and detected using GaAsP photomultiplier tubes (Hamamatsu, H10770PB). Two planes separated by 30 µm within layer 2 were imaged using a Z-piezo. Each plane was acquired at 7.4 Hz with a resolution of 512 × 512 pixels (for rig 1: 1.373 µm/pixel, full FOV 703 µm × 703 µm; for rig 2: 1.367 µm/pixel, full FOV 700 µm × 700 µm). Average laser power at the front of the objective was approximately 70-115 mW. Imaging and behavioral recordings were synchronized as described above.

#### Imaging excitatory neural activity and dopamine dynamics

Simultaneous dual-wavelength imaging was conducted to monitor calcium activity using GCaMP6f and dopamine dynamics using rDA3m. GCaMP6f and rDA3m imaging was performed using the 920-nm fixed-wavelength laser (Axon, Coherent) and 1064-nm fixed -wavelength laser (Axon, Coherent), and fluorescence was collected through a 595/50-nm emission filter. Imaging was performed in layer 2 of the MEC, 100-200 µm below the pial surface, at 30 Hz with a resolution of 512 × 512 pixels (1.116 µm/pixel, full FOV 571.4 µm × 571.4 µm).

#### Imaging calcium activity of excitatory neurons and interneurons

Calcium activity of excitatory neurons expressing jRGECO1a and interneurons (PV+ or SST+) expressing GCaMP8m was imaged in an interleaved manner, during which 1064-nm fixed laser (Axon, Coherent) and 920-nm fixed laser (Axon, Coherent) were applied on alternating imaging frames, respectively, to minimize spectral cross-talk between the two fluorescent signals. Red and green fluorescence was isolated using a 595/50-nm emission filter and a 525/25-nm emission filter, respectively. Imaging was performed in layer 2 of the MEC, 100-200 µm below the pial surface, at 15 Hz with a resolution of 512 × 512 pixels (1.465 µm/pixel, full FOV 750 µm × 750 µm).

### Image data processing

#### Motion correction and ROI identification

In general, imaging data from all experiments underwent cross-correlation-based rigid motion correction as described previously^18^. Regions of interest (ROIs) were identified and fluorescence traces were extracted using Suite2p v0.11.1^91^ with a tau parameter of 0.7 (Ca^2+^ dynamics timescale).

1. Single channel imaging data of GCaMP6 in GP5.3 mice For the multi-environment excitatory-neuron and chemogenetic experiments, ROIs from motion corrected imaging stacks were directly identified using Suite2p.
2. Simultaneous imaging of excitatory neurons and dopamine dynamics Motion correction was first applied to the calcium imaging channel and the shifts of individual calcium frames were applied to the simultaneously acquired dopamine frames so that the two corrected frames were completely aligned. ROIs were identified based on corrected GCaMP6f frames using the same parameters in (1), and were further applied to extract signals in aligned dopamine frames.
3. Interleaved imaging for excitatory neurons and interneurons Motion correction was conducted separately for imaging stacks of excitatory neurons and interneurons. The two corrected imaging stacks were aligned post hoc based on the shift during motion correction. Excitatory neural ROIs from jRGECO1a signals were extracted using Suite2p v0.11.1 with the same parameters as described above.

Since interneuron ROIs could not be properly identified by Suite2p, we conducted a semi-automatic approach. We first used Cellpose-SAM^92^ to automatically detect interneurons from the two-photon imaging recordings. To generate the neuron footprint image for detection, after motion correction the first, middle and the last calcium imaging stacks were concatenated. The mean intensity and standard deviation of each pixel were calculated across frames to obtain the mean intensity projection A and standard deviation projection S. To represent both baseline intensity and fluorescence variation, the neuron footprint image M was generated by multiplying A and S at each pixel. Adaptive histogram equalization was applied to M to enhance contrast between neurons and background. Cellpose-SAM was set up locally in a dedicated Conda environment containing the Cellpose GitHub repository (https://github.com/MouseLand/cellpose/). The detection script of Cellpose-SAM was adapted from the official example notebook (https://github.com/MouseLand/cellpose/blob/main/notebooks/run_Cellpose-SAM.ipynb). The contrast-enhanced footprint image was used as input. The flow_threshold was set to 0.3 and the cellprob_threshold was set to 0.9. Detected interneuron ROIs were then manually curated.

### Fluorescence time course (F) extraction and calculation for ΔF/F and significant transients

1. Excitatory neurons Raw fluorescence was extracted by Suite2p and fractional fluorescence change was calculated as described previously^89^:

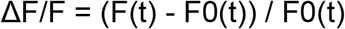 Significant calcium transients were identified using amplitude and duration thresholds selected to produce a 1% false-positive rate^93^. A ΔF/F trace containing only significant calcium transients was used for subsequent analyses unless otherwise stated.
2. Dopamine dynamics Raw fluorescence was directly extracted from motion-corrected imaging stacks and ΔF/F was calculated as described above. Following a previous study^43^, ΔF/F temporal traces were directly used for all analyses without the identification of significant transients.
3. Interneurons For each interneuron ROI, fluorescence time courses were extracted using Suite2p neuropil correction. We adapted the Suite2p pipeline to generate the “stats” data structure containing ROI pixel coordinates and normalized pixel weights (“lam”). Neuropil correction was then performed using Suite2p’s extraction_wrapper function on the motion-corrected imaging data and ROI information to obtain fluorescence traces for downstream analysis. ΔF/F was calculated as described above. Following previous studies^71,94^, ΔF/F temporal traces were directly used for all analyses without the identification of significant transients.

### Data analysis

#### Calcium transient properties

Calcium transient frequency was calculated as the number of significant transients divided by session duration. Mean transient ΔF/F was calculated as the average ΔF/F during significant transients, excluding time points without significant activity. Mean transient duration was calculated as the average duration of significant transients within each session.

To examine the relationship between neuronal activity and calcium transient properties, Pearson correlations were calculated between session-averaged ΔF/F and transient frequency, mean transient ΔF/F, or duration across neurons within each FOV and imaging session. Correlations were calculated separately for each of the 10 FOVs across 15 imaging sessions. P values were corrected using the Benjamini–Hochberg procedure separately across the 150 FOV × session correlations for each transient property, with adjusted P < 0.05 considered significant.

#### Longitudinal cell registration, activity, and relative activity

Neurons across imaging sessions within each FOV were aligned pairwise using a previously developed alignment toolbox^95^. Pairwise alignments were combined across sessions and manually validated and corrected. A neuron was considered active in a session if its ROI was detected by Suite2p.

Session activity was quantified as the session-averaged significant calcium ΔF/F. For analyses of novelty-associated changes in mean ΔF/F, only neurons detected across all imaging sessions were included. Active-neuron counts included all neurons whose ROIs were detected in each session. Changes in mean activity and active-neuron count following novel-environment exposure were calculated as:

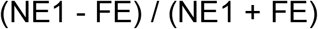

Neuronal persistence was quantified as the number of imaging sessions in which a neuron was detected. To relate novelty-associated neural responses to learning, neural measures were averaged across the two FOVs within each mouse for each novel environment and correlated with predictive licking averaged across days 4-7 of the corresponding novel environment using Pearson correlation. Mouse × novel-environment observations were pooled across the two environments (five mice × two environments, n = 10 observations).

Relative activity was calculated by averaging each neuron’s activity across all sessions in which it was detected and normalizing this value by the mean of these neuron- wise averages within the same FOV. Neurons with relative activity < 1 were classified as low-activity, and neurons with relative activity ≥ 1 as high-activity. For analyses using four relative-activity groups, neurons were divided into ranges of 0-0.5, 0.5-1, 1-1.5, and ≥1.5. Relative activity was calculated using all available imaging sessions in each dataset. For one dopamine mouse in which longitudinal registration was unreliable, activity was normalized within individual sessions instead.

To quantify consistency in relative activity across sessions, Spearman correlations were calculated for each session pair within each FOV using pairwise complete observations. Neurons not detected in one or both sessions were assigned NaN and excluded from the corresponding correlation. We aggregated data by the session pair gap and only visualized session gaps with more than one comparison. Null distributions were generated by independently permuting neuron identities within each session and recalculating the correlations 1000 times. The 95^th^ percentile of the shuffle distribution was calculated separately for each FOV and session pair.

To cross-validate the relative activity classification across sessions, we recalculated relative activity using (1) all sessions except the session being tested, (2) sessions from the other novel environment, or (3) all sessions except the corresponding learning day in both novel environments. For each condition, each neuron’s activity was averaged across the included sessions in which it was detected and normalized by the mean activity across neurons within the FOV. Classification accuracy was calculated as the fraction of neurons assigned to the same low- or high-activity group using the selected sessions and all imaging sessions. Neurons without activity measurements in the selected sessions were excluded from the calculation. Classification accuracy was calculated separately for each FOV and summarized across 10 FOVs from 5 mice.

#### Environment- and learning-day-dependent modeling of neuronal activation

Binary neuronal recruitment data were modeled separately for each FOV using generalized linear mixed-effects models. For each FOV, the neuron-by-day recruitment matrix was converted to long format, with one observation per neuron per day. Neurons were assigned a value of 1 or 0 to indicate whether they were active or inactive, respectively, on a given day. Days 1–7 were assigned to environment B, and days 8–14 to environment C. Learning day within each environment was coded from 1 to 7 and treated as a categorical variable. Logistic mixed-effects models were fitted using MATLAB’s fitglme function. Three candidate models were compared: a null model accounting for neuron-specific differences in overall recruitment probability across all 14 days, an environment model (Env. Model) incorporating environment identity as a two- level categorical predictor (B or C), and a learning model (Learning Model) incorporating learning day as a seven-level categorical predictor. Model fits were compared using the Akaike Information Criterion (AIC)^96^, with lower AIC values indicating a better balance between goodness of fit and model complexity. For each FOV, AIC improvement relative to the null model was calculated by subtracting the AIC of the environment or learning model from that of the null model; positive values indicate improved model support relative to the null model.

In a separate analysis designed to match the number of predictor levels between the environment and learning models, 21 pairwise comparisons of the seven learning days were performed. A separate model was fitted for each learning-day pair, and the resulting AIC improvements relative to the null model were averaged across the 21 pairs for each FOV.

To compare model fits between low- and high-activity neurons, the two populations were downsampled within each FOV to equalize their neuron counts, using the size of the smaller population as the target. For the population with more neurons, downsampling and model fitting were repeated 10 times, and the resulting AIC values were averaged.

#### Spatial activity and spatial field identification

Spatial activity was calculated by binning the significant-transient ΔF/F within 5-cm spatial bins. Time points below the movement-speed threshold were excluded. The speed threshold was determined from a 100-bin histogram of instantaneous positive velocities as twice the center of the first histogram bin, corresponding to approximately 1% of the maximum positive speed.

Significant spatial fields were identified by comparing the spatial activity of each neuron with shuffled responses as described previously^39^. Spatial activity was averaged across laps and smoothed using a 3-bin Gaussian window. For each neuron, 1000 shuffled responses were generated by circularly shifting the temporal ΔF/F trace by a random amount between 0.05 × Nsamples and 0.95 × Nsamples before recalculating spatial activity.

For each spatial bin, the p-value was defined as the fraction of shuffled mean ΔF/F values greater than or equal to the observed value. A spatial field was defined as at least three consecutive bins with activity greater than 80% of shuffled responses at the corresponding positions (1 - p-value ≥ 0.8). Fields at the beginning or end of the track could contain two bins. At least 20% of laps were also required to contain a significant calcium transient within the field. Spatial field coverage was calculated as the proportion of spatial bins for which a neuron possessed field activity. All detected neurons were included, with coverage set to zero for neurons without any significant spatial fields.

Cross-run stability was calculated by correlating each lap’s 5-cm-binned spatial activity with that of every other lap and averaging the valid pairwise correlations.

Spatial information was calculated as described previously^24^:

Spatial information = Σ p_i (f_i / f_mean) log2(f_i / f_mean).

p_i is the fraction of time spent in spatial bin i, f_i is mean significant-transient ΔF/F in that bin, and f_mean is the mean significant-transient ΔF/F across the session. Spatial information was assigned NaN and excluded from analysis when f_mean was zero.

#### Functional cell classification

Cue cells were identified using previously described cue scores^8^. Spatial activity was shifted to obtain the best match to a cue template containing ones at cue locations and zeros elsewhere. Correlations between the shifted activity and cue template were calculated within each cue zone and averaged to obtain the cue score. For each cell, 200 shuffled cue scores were generated by randomizing cue locations. Shuffle distributions were generated separately for each environment by pooling shuffled scores across all cells, FOVs, and imaging days. Cells with cue scores above the 95th percentile of the corresponding shuffle distribution were classified as cue cells.

Grid cells were classified separately on each imaging day using the following criteria^18,39^: (1) at least two spatial fields; (2) more than L/(5w) transitions between in-field and out-of-field periods, where L is track length and w is mean field width; (3) maximum field width <5w; (4) at least 30% of spatial bins assigned to either in-field or out-of-field periods; (5) mean in-field/out-of-field ΔF/F ratio ≥2; and (6) the cell was not classified as a cue cell. If cells satisfied both cue and grid cell criteria, they were classified as cue cells and not grid cells.

Speed scores were calculated as described previously^12^. Position was smoothed with a 0.8-s Gaussian filter before calculating velocity. Time points with velocity <0 or >100 cm/s were excluded. A histogram of speed was used to identify the highest 10-cm/s speed range occupied for at least 30 s, and its mean velocity was used as the upper speed limit. A lower speed limit of 6 cm/s was used. Temporal ΔF/F was smoothed using a 0.4-s Gaussian filter, and speed score was calculated as the Pearson correlation between velocity and ΔF/F within the retained speed range.

For each cell, 1,000 shuffled speed scores were generated by circularly shifting the temporal ΔF/F trace. Shuffle distributions were generated separately for each environment by pooling across all cells, FOVs, and imaging days. Cells above the 99th percentile were classified as positive speed cells and cells below the 1st percentile as negative speed cells.

For each neuron, the fraction of detected sessions in which it was classified as a cue, grid, speed, or unclassified cell was calculated. Speed classification could overlap with cue or grid classification. Cells meeting none of these classifications were considered unclassified for that session.

#### Cross-session / Cross-day spatial stability and remapping

Cross-session or cross-day spatial stability was quantified as the Pearson correlation between the 5-cm-binned spatial activity profiles of the same neuron across two sessions.

For 7 × 7 session correlation matrices, a session × session × neuron correlation tensor was generated separately for each FOV and learning environment. Tensors were concatenated across FOV-learning environment pairs and averaged across neurons.

Adjacent-session stability was calculated within the same environment from the first off-diagonal of session x session neuron correlation tensor. Early and late learning stability were defined as the day 1-day 2 and day 6-day 7 correlations, respectively, and neurons were required to be detected on all four days.

For comparisons between environments of different lengths, the longer environment was restricted to the spatial region corresponding to the shorter environment. For comparisons between 1000-cm and 875-cm tracks, which only occurred on the first novel environment exposure, the last 125 cm of the 1000-cm track was removed.

Remapping between familiar and novel environments was quantified using the same spatial correlation metric, with lower correlations indicating greater remapping.

For learning-dependent single-cell analyses, neurons were required to be detected on NE1 and on at least one of days 5-7. Late-learning values were averaged across the days 5-7 on which each neuron was detected. Changes in within-day consistency and spatial information were calculated as the late-learning value minus NE1.

#### Calcium-event matching

To control for differences in calcium transient properties between low- and high- activity neurons, high-activity calcium transients were adjusted to match the transient frequency, mean transient ΔF/F, and duration of low-activity neurons separately within each FOV and imaging session. The ratio between the mean low- and high-activity values for each transient property within that FOV and session was used for matching and applied to all high-activity neurons in that FOV and session.

Transient number was matched by randomly removing whole high-activity transients, mean transient ΔF/F was matched by scaling transient amplitude, and duration was matched by shortening transients from their end while leaving the rising phase unchanged. All three procedures were applied to the same traces. Spatial activity and significant spatial fields were recalculated after matching, including the 1000 shuffled responses used for spatial field identification. One randomized event-matched dataset was generated for downstream analyses, while relative-activity groups were retained from the original data. To validate the matching procedure, transient frequency, mean transient ΔF/F, and duration were compared between low- and high-activity neurons before and after matching.

#### Spatial field distributions and salient-feature scores

Spatial fields were represented as a binary neuron × spatial-bin matrix, with a value of 1 when a neuron’s spatial field included a given bin. Field distribution was calculated as the fraction of all detected neurons with a field in each spatial bin, including neurons without significant spatial fields in the denominator.

Field distributions were calculated separately for each FOV and learning environment. Early learning was defined as NE day 1, middle learning as the average of NE days 2-4, and late learning as the average of NE days 5-7. Landmark-centered distributions were aligned to the midpoint of each landmark and reward-centered distributions to the reward location. Positions extending beyond the track boundary were padded with NaN.

Landmark and reward salient-feature scores were calculated as:

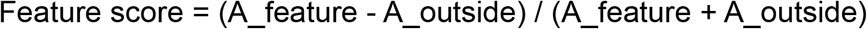

For the main dataset, landmark regions included the landmark itself plus two spatial bins (10 cm) on each side, and the next two bins on each side were used as the outside region. Scores were calculated separately for each landmark and then averaged. The reward region consisted of the reward bin ±6 spatial bins (30 cm), with the next six bins on each side used as the outside region.

For the main excitatory-neuron dataset, feature scores were calculated separately for each imaging day from the population field-distribution trace of each FOV and learning environment, rather than by averaging single-neuron scores. Scores were then averaged across days 2-4 for middle learning and days 5-7 for late learning. The same procedure was used for the four relative-activity groups (0-0.5, 0.5-1, 1-1.5, and ≥1.5). For single- neuron analyses, including feature-preference transitions, A_feature and A_outside were calculated from mean significant-transient spatial ΔF/F. The dopamine and chemogenetic datasets also used single-neuron spatial ΔF/F rather than field distributions to calculate feature scores. When both A_feature and A_outside were zero, the feature score was assigned NaN and excluded from subsequent analyses.

To examine changes in feature preference within the same neurons, cells were classified as reward-preferring when reward score > landmark score + 0.2, landmark- preferring when landmark score > reward score + 0.2, and neutral otherwise. NE1 was used for early classification. Late landmark and reward scores were averaged across days 5-7 on which the cell was detected before classification. Cells were required to be detected on NE1 and at least one late-learning day.

Transition matrices were calculated separately for each FOV × learning- environment pair. Each row was normalized by the number of neurons in the corresponding NE1 starting category. Transition probabilities were compared between low- and high-activity neurons using paired two-sided t-tests across FOV × learning- environment observations, with Benjamini–Hochberg correction across the nine comparisons.

#### Dopamine reward specificity

Dopamine reward specificity was quantified for each neuron using spatially binned dopamine signals (“spatial DA”; 5-cm bins). Pre-reward ramping (R) was defined as the mean spatial DA within the 200 cm preceding the reward location. The non-reward signal (N) was calculated as the mean spatial DA across the remaining track, excluding the 325 cm following reward delivery to minimize contamination from the slow decay of rDA3m fluorescence responses after reward. Dopamine reward specificity was defined as R − N.

#### Comparing calcium reward score or dopamine reward specificity on other days with NE day 1

Neuron-level measurements of the variables were analyzed using a linear mixed- effects model (fitlme function in MATLAB), with day treated as a fixed effect and FOV as a random effect to account for the clustering of neurons within FOVs. The first day in the novel environment (NE day 1) was specified as the reference level, and each remaining day (FE and NE days 2-7) was compared with this NE Day 1 reference. P values were adjusted for multiple comparisons using the Holm-Bonferroni method.

#### Correlations and mediation analysis among dopamine reward specificity, relative activity, and calcium reward score

For each neuron, relative activity, dopamine reward specificity, and calcium reward score were calculated as the mean of the corresponding measure across sessions in which the neuron was active, including the FE and 7 NE days. Pairwise correlations between these measures were calculated across neurons. The correlations between dopamine reward specificity and calcium reward score, and between dopamine reward specificity and relative activity, were compared with null distributions generated from 1,000 circular spatial shuffles of spatial DA. Dopamine reward specificity was recalculated after each shuffle, and the P value was calculated as the proportion of shuffled correlations greater than or equal to the absolute observed correlation.

Mediation analysis^97^ used relative activity as the predictor, dopamine reward specificity as the mediator, and calcium reward score as the outcome. All variables were z-scored within mouse. The indirect effect was calculated as the product of the relative activity–dopamine reward specificity path (a) and reward specificity–calcium reward score path controlling for relative activity (b).

Confidence intervals were estimated using 1,000 hierarchical bootstrap iterations in which mice were resampled with replacement, followed by neurons within each resampled mouse. The 2.5^th^ and 97.5^th^ percentiles of the resulting indirect-effect distribution defined the 95% confidence interval. An indirect effect was considered significant when this interval excluded zero. Percent mediated was calculated as:

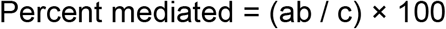

#### Run delay of calcium reward score relative to dopamine reward specificity

This analysis was restricted to low-activity neurons active on both NE days 1 and 2. All 57 neurons from 6 FOVs across 3 mice were pooled for the analysis. The first 16 runs of each imaging session were included, and each measure was smoothed using a rolling average over 4 consecutive runs to reduce noise. Run delay was defined as the relative shift in runs between the calcium reward score and dopamine reward specificity hat maximized their correlation. Positive delays meant that changes in reward score followed changes in dopamine reward specificity.

#### Interneuron and excitatory neuron ROI exclusion based on overlapping level

To reduce fluorescence interference between imaging channels, excitatory neurons and interneurons were excluded if they had any spatial overlap between their ROIs.

#### Excitatory-interneuron coupling and mediation

The PV and SST cohorts contained four and five mice, respectively, with one FOV per mouse and eight imaging sessions (one FE and seven NE sessions). Running speed was regressed separately from each excitatory-neuron and interneuron temporal activity trace using linear regression. Significant-transient ΔF/F was used for excitatory neurons and full ΔF/F for interneurons^71,94^.

For the interneuron datasets, relative activity was calculated across the FE session and seven NE learning sessions. Spatial stability for each excitatory neuron was calculated as the mean of valid pairwise spatial correlations among the seven NE sessions.

For each excitatory neuron, the absolute Pearson correlation between its speed- regressed residual activity and that of each simultaneously imaged interneuron was calculated and averaged across interneurons to obtain its coupling strength. Coupling was calculated separately within each session and averaged across the seven learning sessions for analyses relating coupling to relative activity and spatial stability. For these correlation analyses, neurons were pooled across mice within each PV or SST dataset.

For shuffle controls, each excitatory neuron’s speed-regressed residual trace was independently circularly permuted while interneuron traces remained unchanged. Coupling was recalculated, followed by the relative activity-coupling correlation and the coupling–stability relationship controlling for relative activity. This procedure was repeated 1000 times. Significance was determined the same way as in the dopamine dataset analysis.

Mediation analysis was performed as described above, using relative activity as the predictor, excitatory-interneuron coupling as the mediator, and spatial stability as the outcome.

#### Chemogenetic analyses

Chemogenetic mice were imaged during two FE sessions followed by eight NE learning sessions. Separate cohorts of mice received a single intraperitoneal injection of DCZ (0.5 μg/kg in 50 μL; MedChemExpress, HY-42110) or saline 10 minutes before NE1 imaging. One DCZ mouse was excluded from behavioral and neural analyses because it was substantially older than the remaining cohort. One saline mouse was included in behavioral and remapping analyses but excluded from longitudinal neural analyses because of poor imaging quality from NE4 onward. Treatment assignment was randomized; experimenters were not blinded during treatment administration and imaging. Behavioral analyses used individual mice as the observational units, whereas neural analyses used FOVs (two FOVs per mouse).

Excitatory activity was calculated as the mean significant-transient ΔF/F across neurons detected in all 10 imaging sessions. Delay analyses included only NE1-NE8.

Landmark and reward analyses were performed on low-activity neurons using the salient-feature score described above. For this dataset, landmark regions included the landmark plus four spatial bins (20 cm) on each side, with the next four bins used as the outside region. Reward regions included the reward bin ±7 bins (35 cm), with the next seven bins on each side used as the outside region. Scores were calculated per neuron and averaged within each FOV and day.

Adjacent-day stability was calculated from NE1-NE2 through NE7-NE8. Because longitudinal registration was less reliable in this dataset, only FOVs containing at least 10 neurons detected across all imaging sessions were included. Familiar-to-novel remapping was calculated between FE2 and NE1 using the spatial correlation described above.

Trajectory delays were quantified using the same cross-correlation procedure for behavioral and neural measures. Each DCZ trajectory was cross-correlated with every saline trajectory using integer lags from -3 to +3 days. The lag producing the maximum correlation was identified for each DCZ-saline pair and then averaged across saline comparisons to obtain one delay value per DCZ mouse for behavioral data or per DCZ FOV for neural data. Activity, landmark, reward, and behavioral trajectories contained NE1-NE8, whereas adjacent-day stability contained seven values from NE1-NE2 through NE7-NE8. Positive delays indicated that DCZ trajectories lagged behind saline trajectories. Delay values were compared with zero using two-sided one-sample t-tests.

#### Statistical analysis

Analyses were performed in MATLAB R2024a. Unless otherwise noted, statistical significance was defined as p ≤ 0.05 and tests were two-sided. Paired t-tests were used when the same observational unit contributed to both conditions and unpaired t-tests for independent groups. Pearson correlation was used unless otherwise specified; Spearman correlation was used for activity-rank analyses.

Data are presented as mean ± s.e.m. No formal power calculation was performed; sample sizes were based on typical sample sizes used in the field. Exact sample sizes and experimental units are reported in the corresponding figure legends. Multiple- comparison correction was applied where specified.

---

**Supplementary Figure 1.**
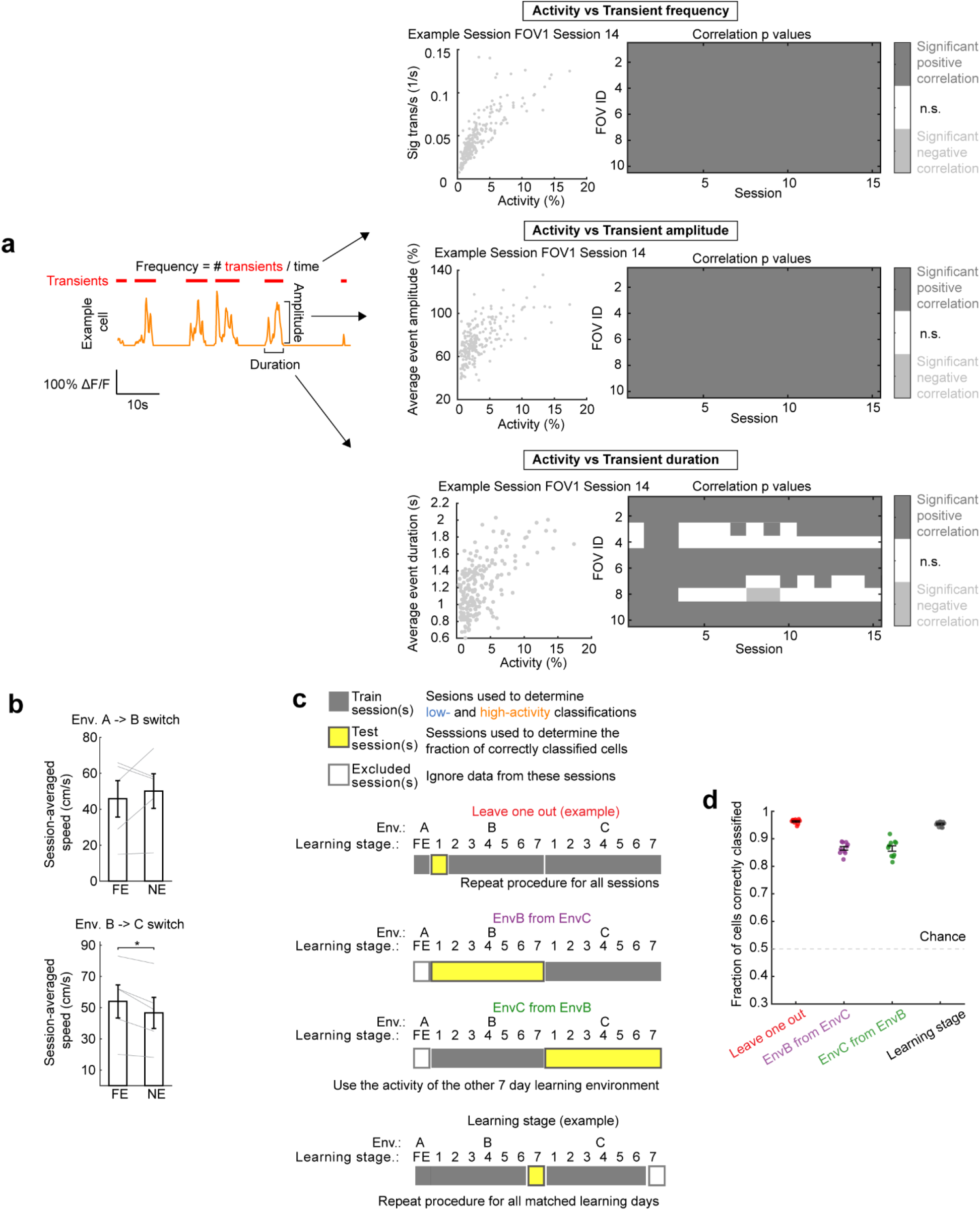
Running speed, calcium transient properties, and cross- validation of relative activity classification. **a.** Relationship between session-averaged neuronal activity and calcium transient properties. Left, representative calcium activity trace illustrating transient frequency, amplitude, and duration. Middle, example Pearson correlations between session- averaged ΔF/F and transient frequency, mean transient ΔF/F, or duration across neurons. Right, significance of correlations across 15 imaging sessions and 10 FOVs from 5 mice. Correlations were calculated separately across neurons within each FOV and session. Dark gray indicates significant positive correlations, light gray indicates significant negative correlations, and white indicates nonsignificant correlations (Benjamini– Hochberg correction across 150 correlations per transient property; adjusted *P* < 0.05). **b.** Mean running speed during FE session and first NE exposure session. Top, environment A exposed to environment B; bottom, environment B exposed to environment C. Running speed was averaged across the entire imaging session, including periods of immobility. Gray lines indicate individual mice (n = 5 mice). **c.** Schematic illustrating cross-validation of classification using all sessions except the session being tested, sessions from the other novel environment, or excluding matched learning days. Yellow indicates test sessions, gray indicates training sessions, and white indicates excluded sessions. **d.** Fraction of correctly classified low- or high-activity neurons within the test sessions using the selected training sessions. Neurons without activity measurements in the test set were not included in the accuracy calculation but were still used to FOV normalize activity. Data are mean ± s.e.m. unless otherwise indicated. The dataset comprised 10 FOVs from 5 mice (2 FOVs per mouse).

**Supplementary Figure 2.**
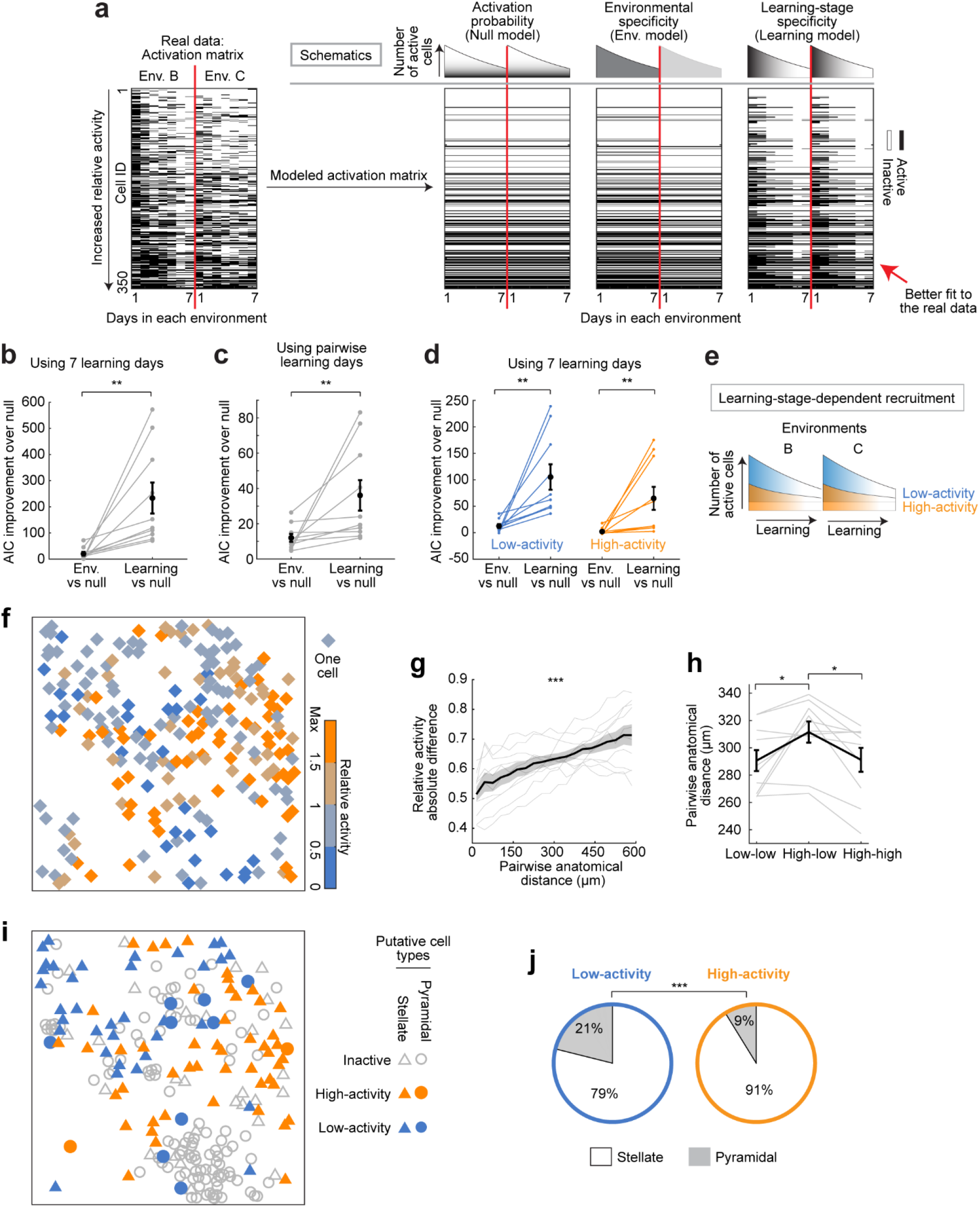
Neural activation dynamics and anatomical organization of relative activity and morphological cell types a-e. We examined whether MEC neural activation dynamics were more strongly associated with environment identity or learning stage. For each FOV, we modeled binary recruitment matrices using a null model that accounted for neuron-specific differences in overall activation probability across days and compared it with models incorporating environment identity (Env. Model) or learning day (Learning Model) as predictors (**a**). Model support was evaluated using the Akaike Information Criterion (AIC)^96^, which balances goodness of fit against model complexity. Across all 10 FOVs, learning day produced a greater improvement in model fit than environment identity (**b**), indicating that cell recruitment was organized predominantly by learning stage. This result persisted when the two predictors were matched for model complexity using pairwise learning-day comparisons, such that both predictors had two levels (**c**). We further examined whether low- and high-activity neurons exhibited different activation dynamics. Learning stage remained the dominant predictor in both populations; however, low-activity neurons showed greater modulation by both learning day and environment (**d**). Thus, low-activity neurons exhibited more intermittent, learning-stage-dependent activation. A similar trend was observed in high-activity neurons, but to a lesser extent, consistent with their more persistent activation (**e**). **a.** Left column: example of neural activation matrix in a representative FOV in tracks B and C, with neurons sorted by their relative activity. Black and white indicate neurons that were active and inactive, respectively, on each session. Remaining columns show modeled activation matrices generated by the null model, Env. model, and Learning model, from left to right. **b.** AIC improvement for the Env. Model and Learning Model relative to the null model, using all seven learning days as predictors. Each line represents one FOV. n=10 FOVs from 5 mice. The sample sizes were the same in **c** and **d**. **c.** AIC improvement for the Env. Model and Learning Model relative to the null model, using all pairwise comparisons between learning days to match the number of predictor levels to that of the Env. model. Results across learning-day pairs were averaged for each FOV. **d.** Same as in **b**, but analyzing low- and high-activity neurons separately. **e.** Schematic illustrating learning-stage-dependent activation of low- and high-activity neurons. **f.** Example FOV showing detected Suite2p ROIs, colored by relative activity level. **g.** Absolute relative activity difference as a function of anatomical distance, pooling all neuron pairs within each FOV. Shading indicates mean ± s.e.m. and gray lines indicate individual FOVs (n = 10 FOVs from 5 mice). **h.** Mean pairwise anatomical distance between low-activity neuron pairs, low-/high- activity neuron pairs, and high-activity neuron pairs. Gray lines indicate individual FOVs (n = 10 FOVs from 5 mice). **i.** Same example FOV as in **f**, showing manually identified putative stellate and pyramidal neurons. Blue and orange indicate low- and high-activity neurons, respectively. Triangles indicate stellate neurons, circles indicate pyramidal neurons, and gray indicates neurons that were manually annotated but not detected as active neurons. **j.** Fraction of putative morphologically classified low-activity and high-activity neurons that were putative stellate or pyramidal. Pie charts show mean proportions across observations (n = 20, 10 FOV × 2 environmental switches from 5 mice). Data are mean ± s.e.m. unless otherwise indicated.

**Supplementary Figure 3.**
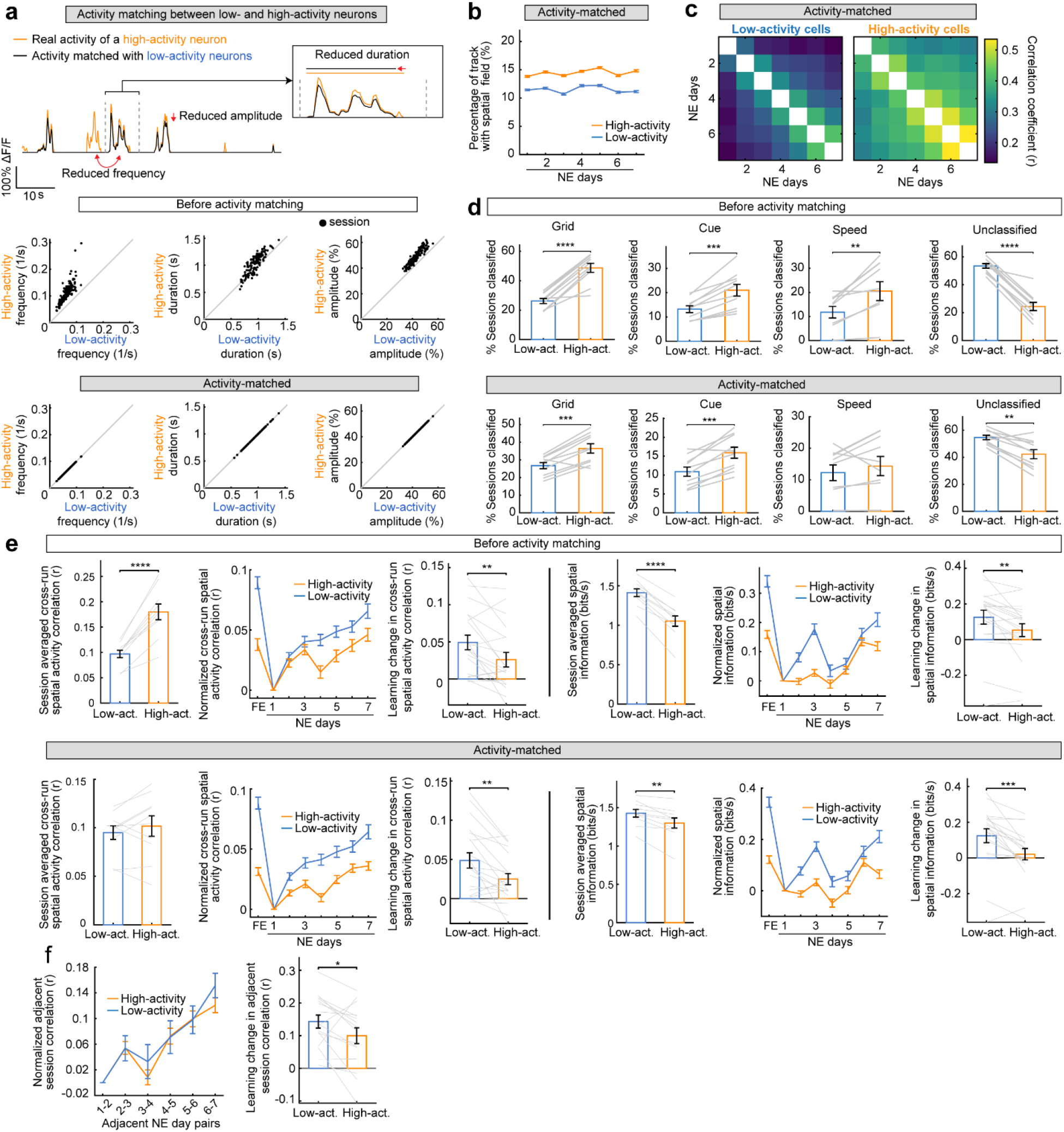
Activity matching and spatial representational properties of low- and high-activity neurons. **a.** Calcium transient activity-matching procedure and validation. Top, representative calcium activity from a high-activity neuron before (orange) and after (black) matching transient frequency, duration, and mean transient ΔF/F to those of low-activity neurons. Bottom, comparisons of transient frequency, duration, and mean transient ΔF/F between low- and high-activity neurons before and after activity matching. Black dots indicate the average from all cells in a session. Solid lines indicate unity. **b.** Spatial field coverage of low- and high-activity neurons after activity matching. Values were pooled across neurons and environments. **c.** Mean pairwise spatial activity correlations across the 7 learning days for low-activity and activity-matched high-activity neurons. Correlations were pooled across neurons. **d.** Percentage of imaging sessions in which low- and high-activity neurons were classified as grid, speed, cue, or unclassified cells, before (top) and after (bottom) activity matching. Gray lines indicate individual FOV observations. **e.** Cross-run stability (left-half) and spatial information (right-half) for low- and high-activity neurons before (top) and after (bottom) activity matching. For each measure, plots show the mean value averaged across all sessions, normalized learning trajectory, and change from NE day 1 to late learning (NE days 5–7). Gray lines indicate individual FOV × learning-environment observations. **f.** Adjacent-session spatial correlations across learning for low-activity and activity- matched high-activity neurons (left), and learning-associated changes in adjacent- session stability from early to late learning (right). Gray lines indicate individual FOV × learning-environment observations. Data are mean ± s.e.m. unless otherwise indicated. The dataset comprised 10 FOVs from 5 mice (2 FOVs per mouse), with 20 FOV × learning-environment observations for the indicated comparisons.

**Supplementary Figure 4.**
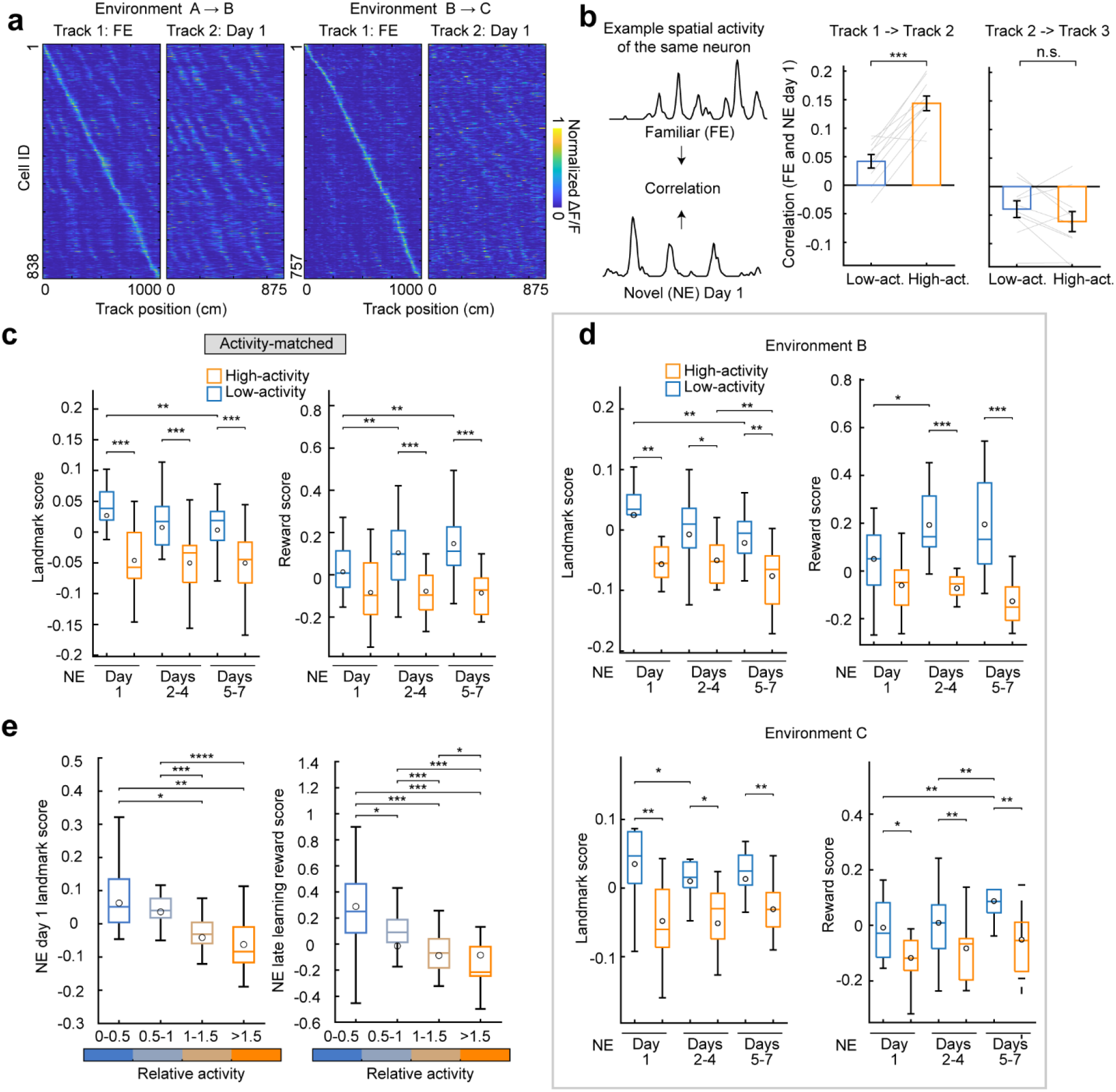
Environmental remapping and feature-specific spatial representations during learning. **a.** Spatial activity of neurons tracked across familiar and novel environments. Neurons are ordered by their peak spatial activity in the familiar environment. **b.** Spatial correlations between familiar and novel environments for low- and high-activity neurons. Left, schematic illustrating the spatial-correlation analysis. Right, mean spatial correlations between environment A and environment B and between environment B and environment C (n = 10 FOVs from 5 mice). **c.** Landmark (left) and reward (right) salient-feature scores for low- and high-activity neurons during NE day 1, days 2–4, and days 5–7 after matching calcium transient frequency, mean transient ΔF/F, and duration in high-activity neurons to those of low- activity neurons (n = 20 FOV × learning-environment observations from 10 FOVs, 5 mice). **d.** Landmark and reward salient-feature scores during learning in novel environments B and C, analyzed separately (n = 10 FOVs from 5 mice per environment). **e.** Landmark scores on NE day 1 (left) and reward scores during late learning (NE days 5–7; right) across four relative-activity groups (0–0.5, 0.5–1, 1–1.5, and ≥1.5) (n = 20 FOV × learning-environment observations from 10 FOVs, 5 mice). Data are mean ± s.e.m. unless otherwise indicated. The dataset comprised 10 FOVs from 5 mice (2 FOVs per mouse). Each FOV × learning environment was treated as one observation for analyses pooling both novel environments and paired t-tests were used to assess significance.

**Supplementary Figure 5.**
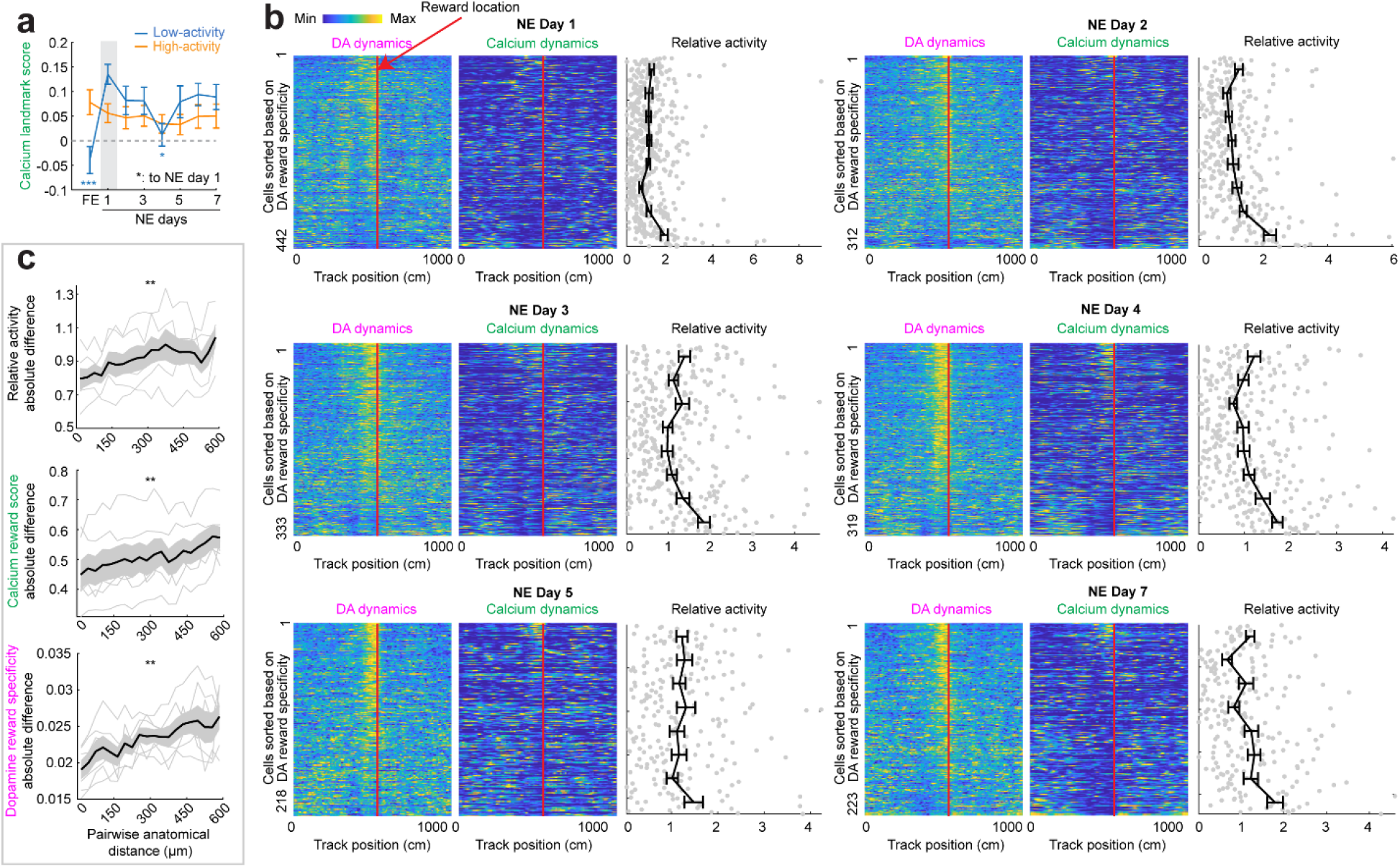
Additional dopamine analysis and anatomical organization of metrics. **a.** Calcium landmark score by activity group. Gray indicates NE day 1, the reference for linear mixed-effects models (day fixed effect, FOV random effect), with Holm-Bonferroni correction. Neurons were pooled across all FOV. **b.** For all NE days, except day 6 (shown in the main figure): DA (left) and calcium (middle) dynamics along the track, and relative activity of individual neurons (right), sorted by DA reward specificity. **c.** Normalized relative activity difference, absolute calcium reward score difference, and dopamine reward specificity difference as a function of pairwise anatomical distance. Grey lines indicate individual FOV. Data are mean ± s.e.m. unless otherwise indicated. The dataset comprised 6 FOVs from 3 mice (2 FOVs per mouse).

**Supplementary Figure 6.**
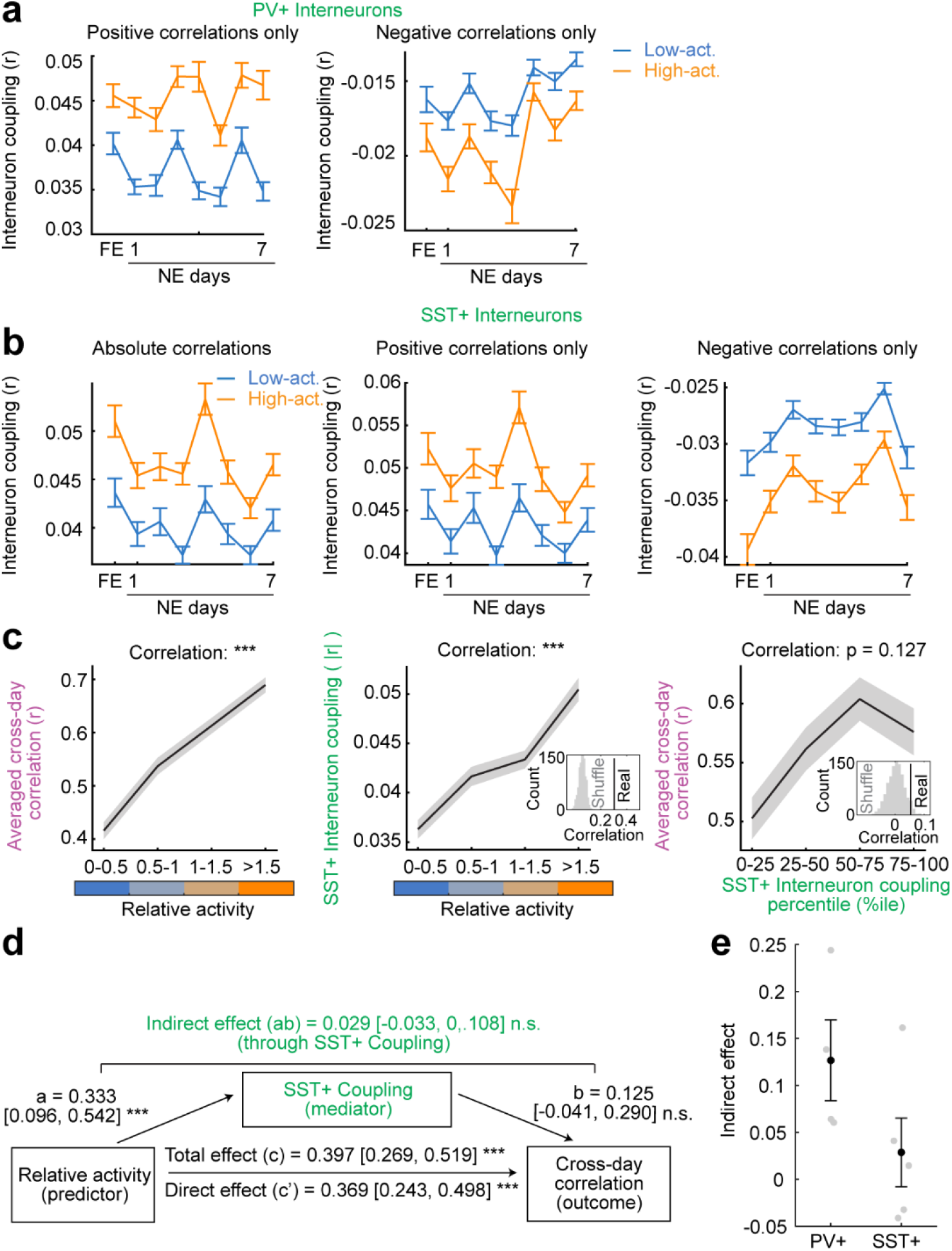
Excitatory–interneuron coupling, representational stability, and mediation in PV+ and SST+ interneuron datasets. **a.** Positive (left) and negative (right) temporal correlations between excitatory neurons and PV+ interneurons across imaging sessions, shown separately for low- and high- activity excitatory neurons. Neurons were pooled across 4 FOVs from 4 mice. **b.** Absolute (left), positive (middle), and negative (right) temporal correlations between excitatory neurons and SST+ interneurons across imaging sessions, shown separately for low- and high-activity excitatory neurons. Neurons were pooled across 5 FOVs from 5 mice. **c.** Relationships among relative activity, SST+ interneuron coupling, and spatial stability. Left, mean cross-day spatial stability across four relative-activity groups. Middle, SST+ interneuron coupling across four relative-activity groups. Right, spatial stability across quartiles of SST+ interneuron coupling. Insets show corresponding shuffle distributions where indicated. The coupling–stability correlation did not reach statistical significance (*P* = 0.127). **d.** Mediation analysis testing whether SST+ interneuron coupling statistically mediates the relationship between relative activity and spatial stability. Total effect of relative activity on spatial stability, 0.397 [95% CI, 0.269, 0.519]; relative activity-to-coupling path, 0.333 [0.096, 0.542]; coupling-to-stability path controlling for relative activity, 0.125 [−0.041, 0.290]; indirect effect through SST+ coupling, 0.029 [−0.033, 0.108]; direct effect after accounting for SST+ coupling, 0.369 [0.243, 0.498]. Confidence intervals were obtained from 1,000 hierarchical bootstrap iterations, resampling mice and then neurons within mice. **e.** Comparison of mouse-level indirect effects through PV+ and SST+ interneuron coupling (PV, n = 4 mice; SST, n = 5 mice). Gray points represent individual mice, and black points and error bars indicate means ± s.e.m. The difference did not reach statistical significance (exact permutation test, *P* = 0.119). Data are mean ± s.e.m. unless otherwise indicated. The PV dataset comprised 4 FOVs from 4 mice, and the SST dataset comprised 5 FOVs from 5 mice (1 FOV per mouse). Running speed was regressed from both excitatory and interneuron temporal activity before calculating correlations. Relative activity, interneuron coupling, and spatial stability were quantified at the single-neuron level. Variables included in the mediation analysis were z-scored within mouse.

**Supplementary Figure 7.**
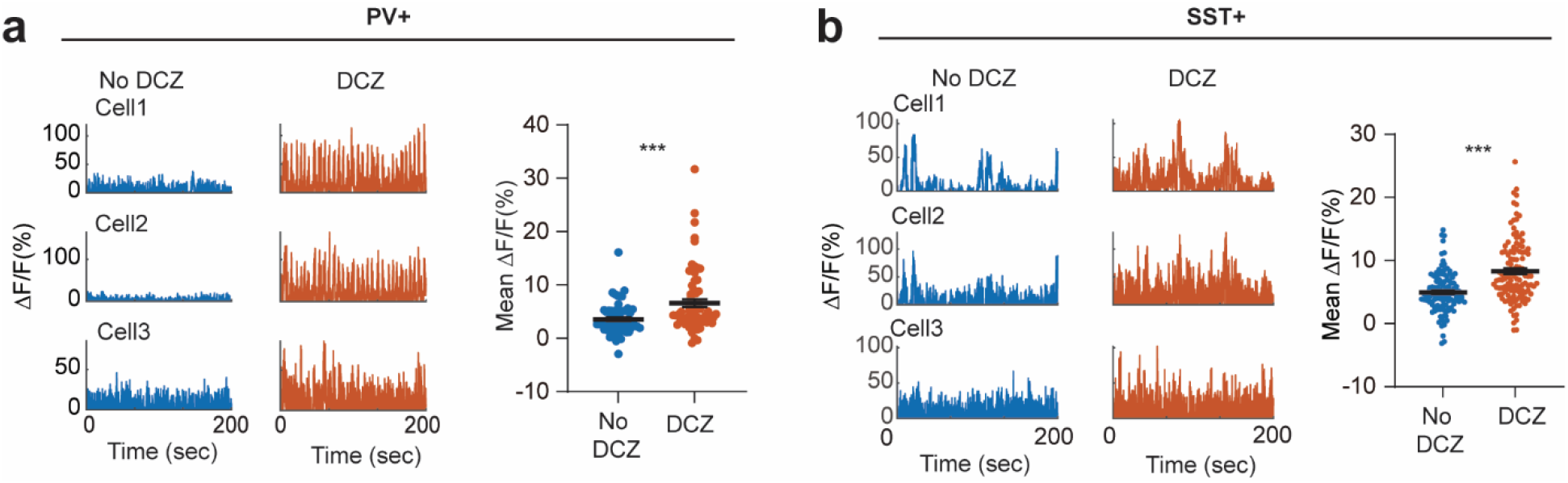
Validation of DCZ-mediated activation of MEC interneurons. **a.** Left, example PV+ interneuron calcium transient activity. Middle, activity traces from the same interneurons after DCZ administration. Right, comparison of mean ΔF/F in PV+ interneurons before and after DCZ administration. **b.** Same as **a** but for SST+ interneurons. Data are mean ± s.e.m. unless otherwise indicated. *, *P* ≤ 0.05, **, P ≤ 0.01, ***, *P* ≤ 0.001; n.s., *P* > 0.05. Neurons were pooled across mice and compared with two-way paired t-tests.

**Supplementary Figure 8.**
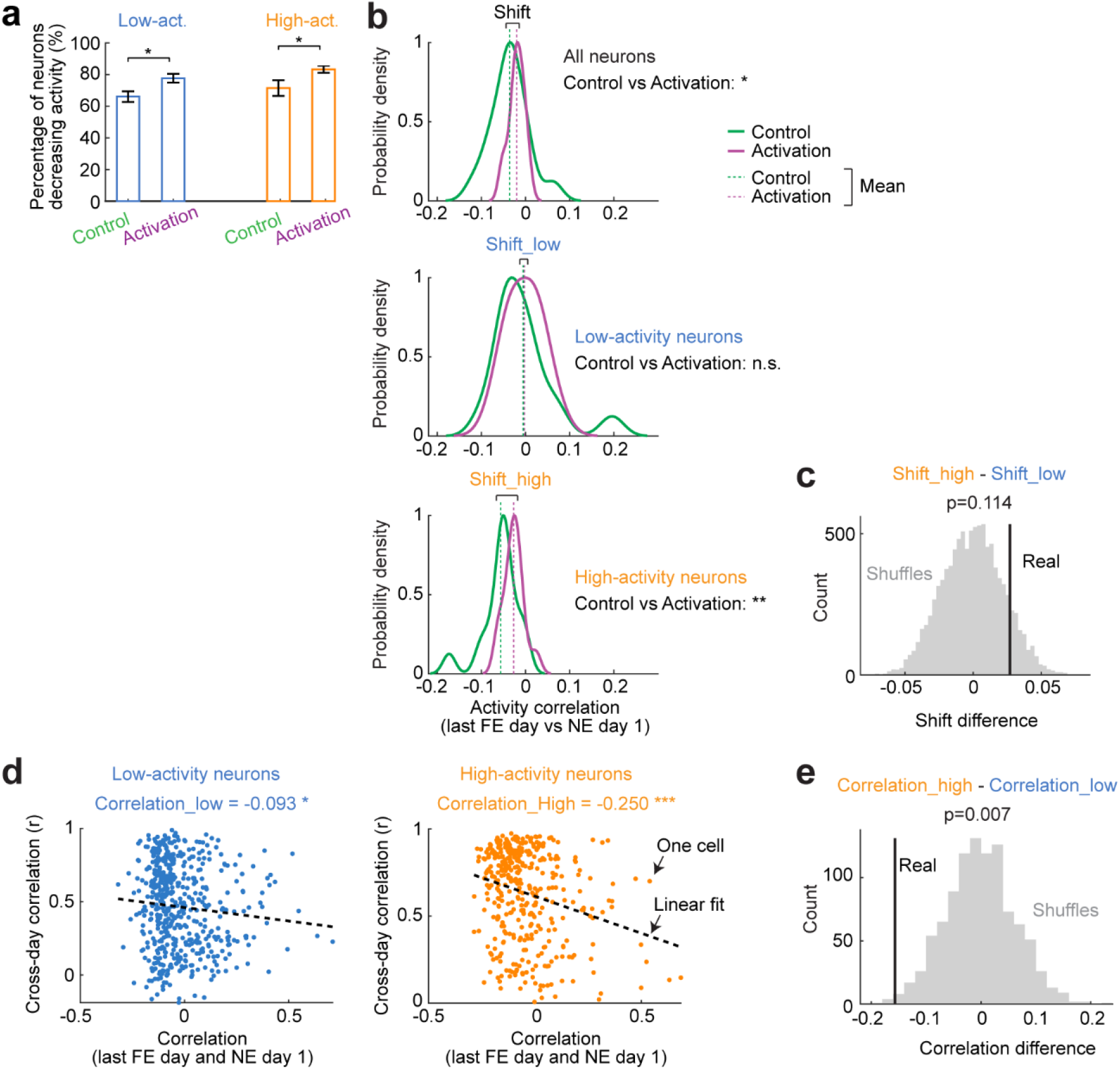
Effects of initial MEC interneuron activation on remapping and representational stability. **a.** Percentage of low- and high-activity neurons that decreased their activity upon exposure to the novel environment in the control and activation conditions. **b.** Kernal density plots comparing spatial activity correlations between the last FE day and NE day 1 in the control and activation conditions across all neurons, low-activity neurons, and high-activity neurons. **c.** Permutation test histogram showing the null distribution of differences in remapping shift between high-activity and low-activity neurons across conditions, generated from 1,000 shuffles. The observed difference is indicated for comparison. The *P* value was calculated as the fraction of shuffled differences greater than the observed difference. **d.** Correlation between remapping correlation (the extent of remapping) and cross-day activity correlation (representational stability during learning) in low- and high-activity neurons of the activation group. Day pairs including NE day 1 were excluded in the cross- day correlation calculation. Neurons were pooled across FOVs. **e.** Permutation test histogram showing null distribution of differences in remapping- stability correlation (as in **d**) between high-activity and low-activity neurons, generated from 1,000 shuffles. The observed difference is indicated for comparison. The *P* value was calculated as the fraction of shuffled differences smaller than the observed difference. Data are mean ± s.e.m. unless otherwise indicated. Analyses in a and b included 16 FOVs from 8 mice in the control group and 14 FOVs from 7 mice in the activation group. Analyses in d and e included 14 FOVs from 7 mice in the activation group.

## Notes

### Competing Interest Statement

The authors have declared no competing interest.

